# Efficient spatial gene expression profiling using split-probe ligation and rolling circle amplification

**DOI:** 10.1101/2025.08.01.668044

**Authors:** Xueqian Xia, Zhaoxiang Xie, Yu Yang, Yanxiu Liu, Weiyan Ma, Bixuan Zhang, Yueping Huang, Yafang Shi, Hui Lin, Lingyu Zhu, Wenhua Li, Chen Lin, Rongqin Ke

**Affiliations:** School of Medicine, Huaqiao University, Xiamen, Fujian 361021, China; Instrumental Analysis Center, Huaqiao University, Xiamen, Fujian 361021, China; Department of Pathology, the 910 Hospital, Quanzhou, Fujian 362000, China

## Abstract

Spatial transcriptomics has transformed our understanding of gene regulation by enabling high-resolution mapping of RNA molecules within their native cellular and tissue environments. This is typically accomplished by capturing or imaging RNA in situ, thereby preserving spatial context. Here, we introduce an *in situ* RNA imaging method based on split-probe ligation and rolling circle amplification (RCA) for profiling spatial gene expression. In this approach, split-probes hybridize to adjacent regions of a target RNA fragment and are then enzymatically ligated to form circular DNA templates, which are subsequently amplified via RCA to boost signal. We demonstrate that this method enables robust *in situ* RNA detection and genotyping in both tissue sections and whole-mount tissue samples. By coupling this technique with *in situ* sequencing, we mapped the spatial expression patterns of 82 genes in the kidneys of healthy and diabetic male and female mice. This analysis revealed distinct localization of *Aqp4* in proximal tubules and principal cells of the collecting ducts, and uncovered sex-specific transcriptomic alterations in diabetic kidneys with spatial resolution.

**GRAPHICAL ABSTRACT:** 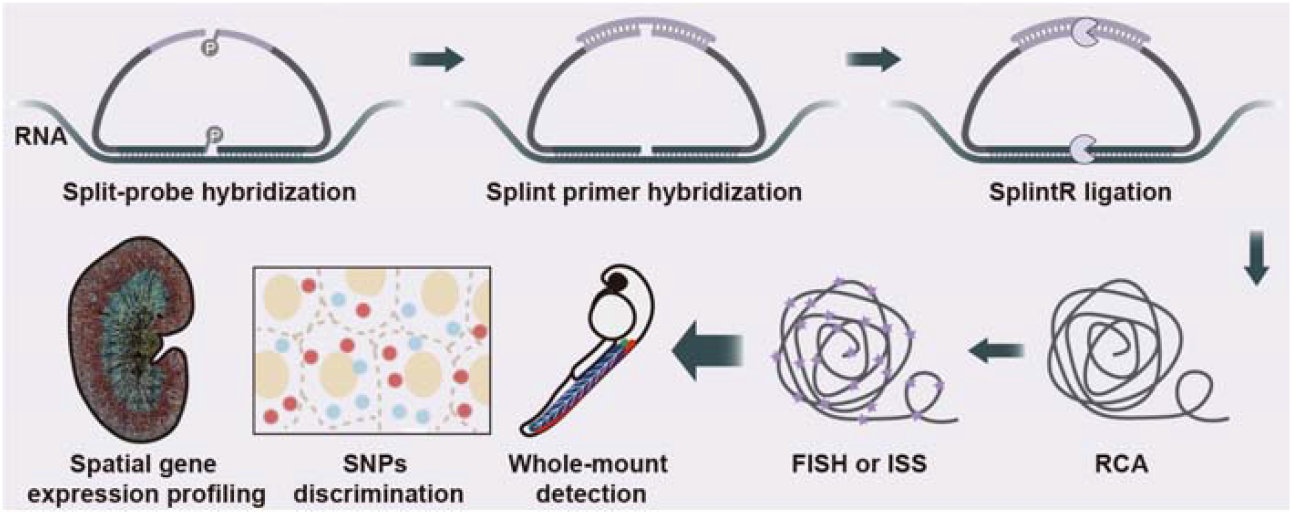

## INTRODUCTION

Profiling the single-cell transcriptome is a powerful approach for investigating the phenotypes and functional diversity of individual cells, providing insights that are unobtainable through bulk analyses. Conventional RNA *in situ* hybridization (ISH) (1,2) has long been used to visualize the spatial distribution of RNA, but is hindered by low sensitivity and specificity, which limit its ability to detect low-abundance transcripts or distinguish closely related sequences. In contrast, single-molecule detection methods have revolutionized transcriptome-level gene profiling by employing iterative imaging cycles to decode gene-specific probes with high precision and accuracy. This advancement relies on amplifying detectable signals by attaching multiple reporters to individual RNA molecules.

Single-molecule fluorescence *in situ* hybridization (smFISH), for instance, utilizes dozens of fluorescently labeled oligonucleotide probes that hybridize to a single mRNA molecule, accumulating sufficient signal for microscopic detection (3,4). This technique set a precedent for visualizing RNA at the single-molecule level *in situ*, laying the foundation for a suite of highly multiplexed RNA detection technologies. Methods like multiplexed error-robust fluorescence *in situ* hybridization (MERFISH) (5) and sequential fluorescence *in situ* hybridization (seqFISH) (6), which achieve transcriptome-scale profiling by combinatorially decoding thousands of RNA species *in situ*, are transforming our understanding of cellular heterogeneity and spatial gene regulation.

Subsequent innovations, such as the hybridization chain reaction (HCR) (7), amplify signals through the triggered polymerization of fluorescent hairpins, thereby enhancing sensitivity without enzymatic reactions. Similarly, branched DNA technology utilizes hierarchical probe assemblies to exponentially increase the number of reporter molecules per target, enabling robust detection (8). Rolling circle amplification (RCA) generates long, repetitive DNA strands from circularized probes, thereby boosting the signal through extensive fluorophore labeling (9). There is a wide range of RCA-based *in situ* RNA profiling techniques (10-12), which are all based on creating circular DNA templates either by circularizing probes through DNA ligation on RNA or cDNA templates, or directly circularizing cDNA itself. Despite being circularized in various ways, padlock probes (13) are classical and the most popular circular DNA precursors for RCA-based RNA detection methods. They can be ligated to cDNA reverse-transcribed from RNA or directly to RNA using different ligases to achieve high efficiency (13).

However, padlock probes are long single-stranded DNA molecules, typically exceeding 60 nucleotides in length, making them relatively expensive to synthesize. To address this limitation, we previously developed a dual-probe ligation strategy for RNA *in situ* hybridization and sequencing (14). This approach conceptually resembles splitting a padlock probe into two shorter oligonucleotides, akin to the principle of *in situ* proximity ligation assay (PLA) (15), which relies on the ligation of two adjacent probes to generate a circular DNA template for rolling circle amplification (RCA). This method, termed amplification-based single-molecule fluorescence in situ hybridization (asmFISH), employs a pair of DNA ligation probes (DLPs) that hybridize adjacent to target RNA sequences. After hybridization, the DLPs are first ligated by a highly efficient DNA ligase that can use RNA as a template. The resulting product is then circularized using a splint DNA oligonucleotide and a second ligase, enabling rolling circle amplification (RCA) for signal enhancement.

While we have demonstrated that asmFISH is highly efficient and robust—successfully detecting diverse RNA species across various fixed cell and tissue types—the use of two sequential ligation steps involving different enzymes introduces redundancy and increases reagent costs. In addition, the method requires a large quantity of Phi29 DNA polymerase for RCA, further elevating assay costs and limiting its scalability and accessibility, especially in resource-limited settings. Here, we present an improved version of asmFISH that addresses the limitations of the original method. Because our approach is based on split-probe ligation, rolling circle amplification, and subsequent fluorescent in situ hybridization, we refer to it as split-roll FISH for simplicity.

In split-roll FISH, the two DNA ligation probes are both ligated and circularized in a single step using SplintR Ligase, a DNA ligase derived from Chlorella virus PBCV-1 that efficiently joins adjacent DNA oligonucleotides hybridized to an RNA template (16). To reduce enzyme consumption during amplification, we replaced conventional Phi29 DNA polymerase with EquiPhi29, a thermostable variant with higher processivity and yield, enabling efficient rolling circle amplification (RCA) using substantially lower enzyme amounts. Additionally, we optimized the reaction protocols and increased the incubation temperatures to further enhance enzymatic reaction efficiency.

We applied split-roll FISH to fresh frozen (FF), formalin-fixed paraffin-embedded (FFPE), and whole-mount tissue samples. Compared to the previous whole-mount protocol for zebrafish larvae, our improved method reduced the total assay time from 25 to 33 hours to just 9 hours. Furthermore, we established a strategy for detecting single-nucleotide polymorphisms (SNPs), enabling sensitive detection of SNPs in both cells and tissues. By integrating split-roll FISH with an improved in situ sequencing (ISS) protocol (17), we developed split-roll ISS, a split-probe ligation and RCA-based in situ sequencing method. Using this approach, we profiled the expression of 82 genes in mouse kidneys. Spatial gene mapping in healthy and diabetic male and female mice revealed that *Aqp4* is predominantly expressed in the proximal tubules and principal cells of the collecting ducts.

Overall, our study demonstrates that the split-probe ligation and RCA-based method enables fast, accurate, and cost-effective multiplexed RNA in situ detection.

## MATERIAL AND METHODS

### Probe design and phosphorylation

The genes detected in the whole-mount zebrafish tissue experiment were manually curated based on previously published literature (7,18), and the sequences are listed in Supplementary Table S1. For the mouse kidney experiments, genes were selected based on publicly available single-cell RNA sequencing (scRNA-seq) datasets (19,20), supplemented with differentially expressed genes (DEGs) identified from bulk RNA-seq, and the sequences are listed in Supplementary Table S2.

Split probes for *in situ* detection have been previously described (14). After ligation and circularization, they contain a 32-nucleotide region complementary to the target mRNA and a sequence identical to the detection probe. Split-probes for in situ sequencing (ISS), as previously reported (17), retain these features and additionally incorporate barcode sequences that are specifically recognized by interrogation probes (Supplementary Table S2-3).

For SNP genotyping assays, the upstream and downstream probes each contain a sequence identical to the detection probe. The downstream probe also includes a region complementary to the target mRNA, which is designed to span the polymorphic site. Distinct detection probe sequences are assigned according to the specific nucleotide variant. All probe sequences are listed in Supplementary Table S4.

All split-probes were phosphorylated before use. A reaction mixture containing 0.2 U/mL T4 polynucleotide kinase (T4 PNK; Thermo Fisher Scientific, EK0032), 1× PNK Buffer A, 1 mM ATP (Thermo Fisher Scientific, R0441), and 2 µM split-probes (Sangon Biotech, China) was incubated at 37⍰°C for 30 minutes, followed by heat inactivation at 65⍰°C for 10 minutes. The phosphorylated probe mix was then stored at −20 °C until use.

### Cell culture and sample preparation

SW480 (ATCC) and NCI-H157 (National Collection of Authenticated Cell Cultures, China) cells were cultured in RPMI-1640 (BasalMedia, L230KJ), SKOV3 (The Cell Resource Sharing Platform of the National Laboratory, China) were cultured in McCoy’s 5A Medium (BasalMedia, L630KJ), while MCF-7 (ATCC) and A549 (ATCC) cells were cultured in DMEM (Sangon Biotech, E600003). All culture media were supplemented with 10% fetal bovine serum (Gibco, 10099-141C) and 1% penicillin-streptomycin solution (BasalMedia, S110JV). For NCI-H157 cells, the medium was additionally supplemented with 10 mM HEPES (Sangon Biotech, E607018) and 1 mM sodium pyruvate (BasalMedia, S410JV). Cells were incubated at 37^°^C in 5% CO_2_.

To prepare cell culture slides, cells were detached from the culture flask with 0.25% (w/v) trypsin-EDTA and seeded on adhesive glass slides (Citotest, 188105) submerged in the corresponding culture medium in a 150 mm diameter Petri dish (Jet Biofil, TCD110150). The cells were then allowed to grow until 70% confluence, and then washed three times with diethyl pyrocarbonate (Sangon Biotech, B600154) treated 1× phosphate-buffered saline (1× DEPC-PBS) after removal of the culture medium from the Petri dish. Next, fixation was performed by incubating the slides in 4% paraformaldehyde (PFA) (Sigma-Aldrich, 16005) in 1× DEPC-PBS for 30 min at room temperature. After discarding the PFA fixative, the cells were washed twice for 3 min each in 1× DEPC-PBS, followed by dehydration in an ethanol series of 70%, 85%, and absolute for 5 min each. After air-drying at room temperature, slides were stored at −80°C until use. Before use, the adherent cell samples were permeabilized with 0.1 M HCl for 5 min and washed twice with 1× DEPC-PBS supplemented with 0.1% Tween-20 (DEPC-PBST).

### Pre-treatment of FFPE tissue samples

Human colorectal cancer FFPE tissue sections were obtained from the Department of Pathology, the 910th Hospital, Quanzhou, Fujian, China. The 9-week-old *LSL*-*Kras*^G12D/+^; *Sftpc*-Cre mice (hereafter referred to as KS) and *LSL*-*Kras*^G12D/+^ mice (hereafter referred to as WT) lung FFPE tissue sections were provided by Xiamen University. Details regarding the KS and WT mice have been described previously (21). FFPE tissue sections were first baked at 60°C for 30 min and then dewaxed in xylene twice for 15 min and 10 min, respectively. The rehydration process was then completed by submerging the sample in absolute ethanol, 95% ethanol, and 70% ethanol for 2 minutes each, followed by dipping in DEPC-treated H_2_O (DEPC-H_2_O) for 5 minutes and washing with 1× DEPC-PBS for 2 minutes. The tissue sections were fixed with 4% PFA in 1× DEPC-PBS for 10 min at room temperature and then washed with 1× DEPC-PBS for 2 min. Next, a solution of 0.1 M HCl containing 0.3 mg/mL pepsin (Sigma-Aldrich, P7012) was applied to the tissue sections for 30 min at 37°C. After washing with DEPC-H_2_O for 5 min and DEPC-PBS for 2 min.

### Pre-treatment of FF mouse kidney tissue samples

BKS-*Lepr*^em2Cd479^/Gpt (hereafter referred to as db/db) mice (Strain NO. T002407) and C57BLKS/JGpt mice (background mice, db/m) (Strain NO. N000214) were purchased from GemPharmatech (Nanjing, China). In Week 13, mice were anaesthetized through an intraperitoneal injection of tribromoethanol (4 mg/10 g), followed by transcardiac perfusion with saline, and then both kidneys were dissected. One of the kidneys was taken for bulk RNA-seq and the other was embedded in optimal cutting temperature (OCT, SAKURA, 4583) compound and frozen at −80°C. The frozen OCT blocks were trimmed and sectioned at a thickness of 10 µm onto adhesive glass slides using a Leica cryostat (Leica, Germany). The slides can be stored at −80°C until used. Before use, FF tissue sections were first fixed in 4% PFA in 1× DEPC-PBS for 10 min, followed by washing in 1× DEPC-PBS for 2 min twice. The slides were then incubated in 0.1 M HCl for 10 min. Finally, three washes were performed in 1× DEPC-PBST.

### Pre-treatment of zebrafish embryo samples

The wild-type adult zebrafish (Danio rerio) were raised in circulating water at 28.5 ± 0.5 °C, and pH 7.0 ± 0.25. They were fed three times a day with saltwater shrimp that had been incubated for more than 24 h. Adult male and female zebrafish were placed in a mating box at a 4:1 ratio, separated overnight with a partition. The partition was removed during the next photoperiod to allow mating, and embryos were collected. 27 h post-fertilization (27 hpf) embryos were collected into a 1.5 mL Eppendorf tube, and excess egg H_2_O was removed. Embryos were then fixed in 4% PFA for 24 hours at 4°C. The embryos were then washed three times for 5 minutes in DEPC-PBS. Dehydrating and permeabilizing were carried out with four 10-minute 100% methanol washes. Embryos can be stored at −20°C until used. Before use, embryos were rehydrated with a series of 75%, 50%, and 25% methanol in DEPC-PBST, 100% DEPC-PBST washes, 5 times for 5 min each. Then, embryos were permeabilized with 0.2 M HCl for 10 min and subjected to proteinase K (Sangon Biotech, B600452; 5 μg/mL in PBS) for 5 min at room temperature (RT). After being fixed again with 4% PFA for 5 min, the embryos were washed three times with PBST. Then, pre-hybridizing with 150 μL of hybridization buffer (6× SSC, 10% formamide) for 30 min at 42^°^C.

### Split-probe hybridization and circularization

A mix containing 0.1 μM phosphorylated split-probes (for *in situ* sequencing in mouse kidneys, containing 0.02 μM phosphorylated split-probes), 6× SSC, 5% formamide and 5% PEG4000 was added to the sample for 4 h at 42°C. Then, wash the reaction area with DEPC-PBST thrice. Splint oligonucleotides were hybridized in a mixture containing 0.5 μM of splint oligonucleotide (Supplementary Table 1, 2 and 4), 6× SSC, 10% formamide, and 5% PEG 4000. The mixture was incubated for 30 min at 42°C, followed by three washes with DEPC-PBST. The two nicks in the split-probe were ligated to form a circle by SplintR Ligase using mRNA and splint oligonucleotide as templates. The ligation reaction mixture contained 0.5 U/μL SplintR Ligase (NEB, M0375L), 1× SplintR reaction buffer, 15% glycerol (Sigma Aldrich, G5516), 0.2 μg/μL BSA (Sangon Biotech, B600036), and 1 U/μL RiboLock RNase inhibitor (Thermo Fisher Scientific, EO0384). The mixture was incubated at 42°C for 30 min, followed by three washes with DEPC-PBST.

### Hybridization and circularization of asmFISH probes

For asmFISH, the protocol differed from the above at the splint oligonucleotide hybridization and ligation steps, while the probe hybridization step remained consistent. After probe hybridization, DLP ligation was carried out by adding a reaction mix containing 0.5 U/μL SplintR Ligase, 1× SplintR reaction buffer, 50% glycerol, 0.2 μg/μL BSA, and 1 U/μL RiboLock RNase inhibitor, followed by incubation at 37°C for 30 min. Subsequently, the ligated probes were circularized using a T4 ligase-mediated DNA ligation mix, which included 0.1 U/μL T4 DNA ligase (Thermo Fisher Scientific, EL0012), 0.5 μM of splint oligonucleotide (Supplementary Material-2), 0.2 μg/μL BSA, and 1 U/μL RiboLock RNase inhibitor in 1× T4 DNA ligase reaction buffer, with incubation at 37^°^C for 30 min.

### Digestion of the excess single-stranded probe

This step was only conducted in the zebrafish embryo whole tissue ISH assay. The reaction mixture contained 0.4 U/μL Exonuclease I (Thermo Fisher Scientific, EN0581), 1× Exonuclease I reaction buffer, 5% glycerol, and 0.2 μg/μL BSA. The mixture was incubated at 37°C for 30 min, followed by three washes with DEPC-PBST.

### RCA of split-roll FISH

RCA was initiated by adding a reaction mixture in DEPC-H_2_O containing 1× EquiPhi29 DNA polymerase reaction buffer, 5% glycerol, 0.2 μg/μL BSA, 1 mM dNTPs (Thermo Fisher Scientific, R0182), 0.2 U/μL EquiPhi29 DNA polymerase (Thermo Fisher Scientific, A39392), 1 mM DTT and 5% PEG 4000 to the chamber and incubated 2 h (cells) or 3 h (tissue samples) at 42^°^C, followed by three washes with DEPC-PBST. To ensure that the staff have enough rest, the RCA can be left overnight at 30°C, and the experiment can be continued the next day. This will not affect the results.

### RCA of asmFISH

RCA was initiated by adding a reaction mixture in DEPC-H_2_O containing 1× Phi29 DNA polymerase reaction buffer, 5% glycerol, 0.2 μg/μL BSA, 1 mM dNTPs, 1 U/μL Phi29 DNA polymerase (Thermo Fisher Scientific, EP0094), to the chamber and incubated 2 h for cell or 3 h for tissue samples at 37^°^C, followed by three washes with DEPC-PBST. The RCA can be left overnight at 30°C, and the experiment can be continued the next day. This will not affect the results.

### Detection of RCA products (RCPs) by FISH

0.1 μM fluorescently labeled detection probes (Supplementary Table 1 and 4) in 2× SSC and 20% formamide were applied and incubated for 30 min at 42°C. Samples were then washed three times with DEPC-PBST. For cells or thin tissue samples, SlowFade Gold Antifade Mountant medium (Invitrogen, S36936), containing 0.5 μg/mL DAPI for nuclear staining, was applied to the slides for mounting. Then, the images were acquired using a Leica DM6B fluorescence microscope (Leica, Germany) equipped with either a DFC9000GT (Leica, Germany) camera or a K8 camera (Leica, Germany), both of which were used with a 20× objective (Leica, Germany). For mounting the zebrafish embryo, a chamber was created by aligning two 20 mm × 20 mm coverslips apart on a 25 mm × 75 mm glass slide. A hydrophobic circle was created using an ImmEdge Pen, and then the embryo was transferred to the circle. Approximately 30 μL of mounting medium containing 0.5 μg/mL DAPI was added to the slide, and the embryos were placed on the medium, oriented for lateral imaging. A 20 mm × 50 mm coverslip was placed on top of two glass slides to close the chamber. Samples were visualized and imaged with a Leica SP8 confocal microscope.

### Detection of RCPs by improved *in situ* sequencing

A mixture containing 0.2 μM anchor primer (Sangon Biotech), 1× T4 DNA ligase buffer, 0.1 U/mL T4 DNA ligase, 0.2 μM fluorescently labeled sequencing interrogation probes (Invitrogen), 2 mM ATP was applied and incubated at 30°C for 45 min, followed by washing three times with 1× DEPC-PBST, and air-dry. The slides were then mounted in SlowFade Gold Antifade Mountant medium containing 0.5 μg/mL DAPI. Then the images were acquired by a Leica DM6B fluorescence microscope equipped with a DFC9000GT camera (Leica) using a 20× objective (Leica). To strip off ligated sequencing probes and prepare the slides for the next sequencing cycle, the slides were treated two times with 2× SSC at RT for 10 min each, three times with stripping buffer that contained 80% formamide and 0.1% Triton X-100 at 37°C for 10 min each, and washed three times with PBST. The same procedures were applied for each sequencing cycle by repeating the hybridization and ligation of the sequencing probes, imaging, and stripping off. To enhance detection throughput, we employed two splint primers and their corresponding two sets of anchor primers. Theoretically, this approach enables batch-wise sequencing across 8 rounds, allowing the detection of 512 genes.

### Image sequencing data pre-processing

For the fluorescence images generated after *in situ* hybridization, we first used *filters*.gaussian in the skimage package to smooth the images, and *blob_log* was used to detect blobs in the images. The fluorescence intensity was segmented between the signal and the background using *threshold_otsu* in the skimage package (22).

For the microscopy data after improved *in situ* sequencing, we first find matching points between different rounds based on the DAPI channel of the reference round, create a transformation matrix, and calibrate other channels to perform registration. Subsequently, spot detection is implemented in the enhanced image after top-hat filtering. The fluorescent signals in different rounds are connected into barcodes and matched with the corresponding gene barcodes. These steps are performed by a common script in the laboratory.

### Cell segmentation

To create the single-cell expression matrix, we first used cellpose3 to identify the nucleus outlines of the grayscale DAPI images of the reference round. We used the latest “cyto3” model and set the *flow_threshold* to 0.8 (23). To ensure that the RCP in the cytoplasm is assigned to the cell, we expanded the original mask by 20 pixels. Finally, the RCPs were assigned to the nearest cell. The visualization results were achieved using *segmentation. find_bondaries* in skimage and *pyplot*.*scatter* in matplotlib.

### Single-cell spatial analysis

The cell type annotation results were defined using the expression ratio of cell marker genes in each cell. For each cell, the values of the original expression matrix were converted to percentage values and the sum of the probabilities of each cell type gene set was calculated. Each cell was labeled as the cell type with the highest score (17).

We used the labels of zonation structure in the single-cell data GSE129798 to map the mouse kidney *in situ* sequencing data. After applying *pp_adatas* in the Python package Tangram to preprocess the single-cell data and spatial data, we used the *map_cells_to_space* function to achieve the mapping (20,24). For neighborhood enrichment analysis, we used the squidpy function *gr*.*spatial_neighbors* to calculate the neighbor graph, and then we could use *gr*.*nhood_enrichment* to calculate the scores between different cell types and visualize them using *pl*.*nhood_*enrichment. The function *gr*.*spatial_autocorr* was used to calculate the Moran’s I score of differentially expressed genes (25).

Finally, we created a Seurat object for the mouse kidney data and imported the corresponding cell type annotation results and sample information. After integrating the 8 samples, we used *AverageExpression* to obtain the average expression of each sample. After inputting the above mean matrix into the *gsva* function, we used KEGG gene sets (MSigDB, C2) to perform correlation analysis and draw a heat map. KEGG enrichment analysis of differentially expressed genes was implemented using the R package clusterprofilter (26-28).

### Ethical statement

The use of archival human-derived tissue sections was approved by the Ethics Committee of the School of Medicine, Huaqiao University. For animal studies, all experimental procedures were performed in strict accordance with *the Guidelines for the Care and Use of Laboratory Animals* and other relevant ethical regulations, and all protocols were approved by the Ethics Committee of the School of Medicine, Huaqiao University, with specific approval numbers: zebrafish (A2021046), mice (A2024036, A2024051).

## RESULTS

### Implementation of split-roll FISH

The split-roll FISH is developed from our previously published asmFISH method. As illustrated in Figure 1A, the split probes are adjacently hybridized to target RNA fragments, followed by the hybridization of splint oligonucleotides on each pair of split probes, forming circular DNA structures with two nicks. Subsequently, the two nicks are sealed by SplintR DNA Ligase, which can mediate DNA ligation using either DNA or RNA as templates, resulting in complete DNA circles. RCA is then carried out using the splint oligonucleotides as primers by EquiPhi29 DNA polymerase, a thermostable variant of the original Phi29 DNA polymerase. The RCA products (RCPs) can then be detected either by FISH or ISS, depending on the number of genes analyzed. Hybridizing spectrally distinct fluorescently labeled detection probes with the RCPs enables the detection of multiple genes in a single imaging cycle. While using ISS to encode and decode the RCPs, hundreds of genes can be detected through multiple imaging cycles.

**Figure 1:**
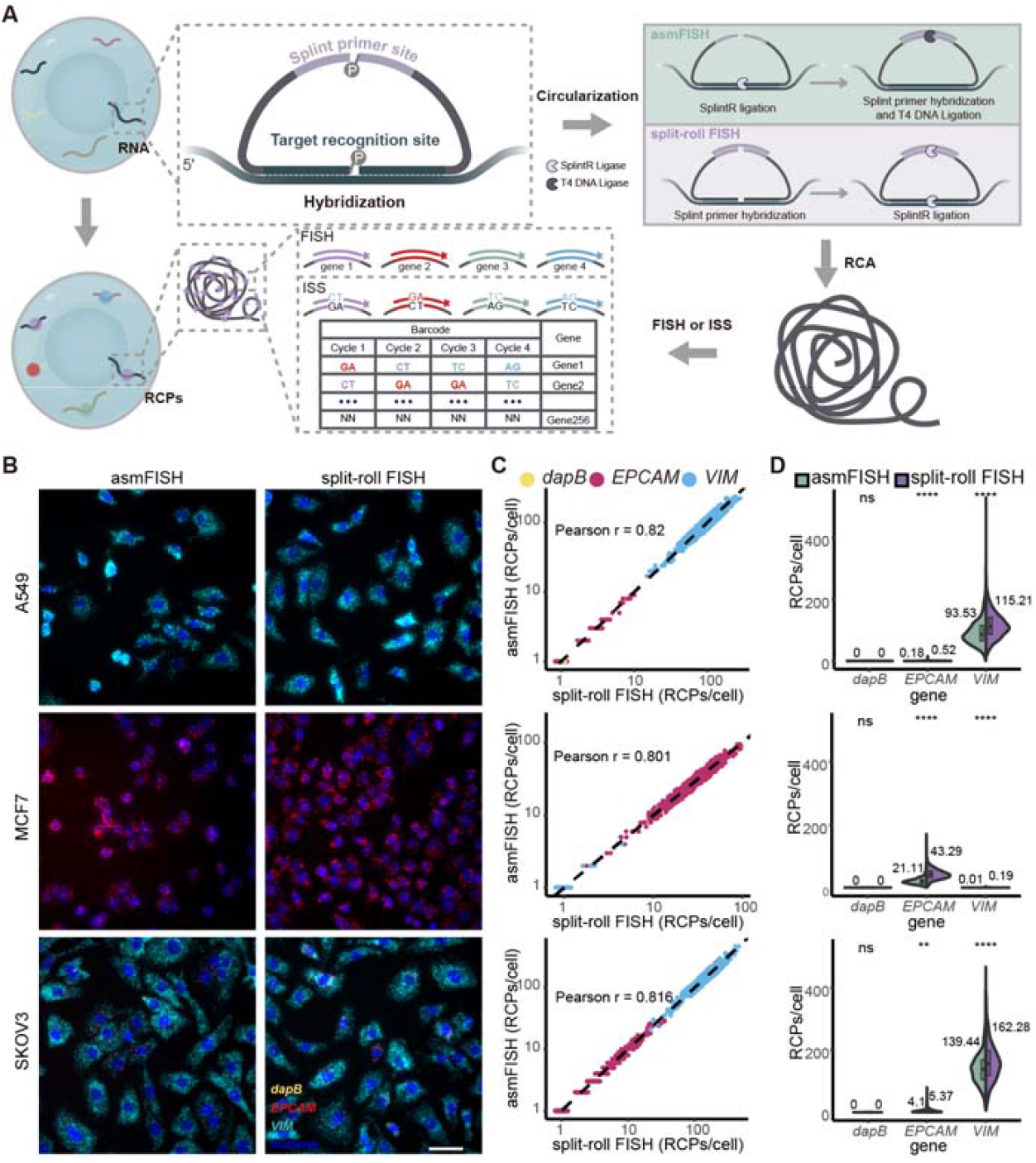
Comparison of split-roll FISH and asmFISH. (**A**) Schematic diagram of split-roll FISH and asmFISH. In split-roll FISH (purple frame), split-probe ligation and circularization were carried out simultaneously by SplintR Ligase only, while in asmFISH (green frame), SplintR DNA Ligase was used to ligate split probes first, followed by using T4 DNA Ligase for subsequent probe circularization. (**B**) Comparison of the detection efficiency of split-roll FISH and asmFISH in different cells. Scale bar: 50 μm. (**C**) Comparison of the measurement results of split-roll FISH on different cell markers with asmFISH. (**D**) Violin plots of the detected RCPs per cell for the corresponding genes in different cells. Statistical significance between methods was assessed using two-sided unpaired t-tests. (****) indicating *P* < 0.0001. (**) indicating *P* < 0.01. “ns” denotes no significance (*P* > 0.05).

We compared the detection efficiency of the split-roll FISH to asmFISH by detecting different transcripts in cell cultures. The bacterial gene *dapB* from Bacillus subtilis strain SMY was used as a negative control, while *EPCAM* and *VIM*, representing different expression levels, served as targets for comparison. All probe sequences are listed in Supplementary Table S1. Then, the detection of *EPCAM, VIM*, and *dapB* mRNA expression in A549, MCF7, and SKOV3 cells was compared using asmFISH and split-roll FISH (Figure 1B). Both assays showed 0 RCPs from *dapB* in all cell cultures, indicating that split-roll FISH exhibits high specificity consistency with asmFISH (Figure 1C). However, the split-roll FISH had a higher detection efficiency than asmFISH, as illustrated by the detection of more RCPs from *EPCAM* and *VIM* in all cell lines (Figure 1D). For A549 cells, the mean number of RCPs per cell detected by asmFISH was 0.18 (n = 1,122) for *EPCAM* and 93.53 (n = 1,122) for *VIM*; for split-roll FISH, the corresponding values were 0.52 (n = 1,068) for *EPCAM* and 115.21 (n = 1,068) for *VIM*. In MCF-7 cells, asmFISH yielded a mean of 21.11 (n = 1,379) RCPs per cell for *EPCAM* and 0.01 (n = 1,379) for *VIM*, whereas split-roll FISH detected 43.29 (n = 1,253) RCPs for *EPCAM* and 0.19 (n = 1,253) for *VIM*. In SKOV3 cells, asmFISH showed a mean of 4.1 (n = 983) RCPs per cell for *EPCAM* and 139.44 (n = 983) for *VIM*, while split roll FISH detected 5.37 (n = 1,045) RCPs for *EPCAM* and 162.28 (n = 1,045) for VIM. These results demonstrate that both methods are consistent in target detection across genes with varying expression levels, while split-roll FISH offers improved sensitivity. To further improve the detection efficiency of the new method, we optimized the components of the split probe hybridization and circularization system (Supplementary Fig. S1). Based on the results of the assay, we selected 5% formamide and 6× SSC as the optimal conditions for split-probe hybridization and 15% glycerol as the optimal concentration for split-probe circularization. We then used these optimized conditions in subsequent experiments.

### RNA detection on human FFPE colon cancer tissue sections

To show that split-roll FISH can be applied to clinically relevant samples, we tested the detection of the same genes (*dapB, EPCAM, VIM*) in human FFPE colon cancer tissue sections. *EPCAM*, a classic cancer marker, is of great significance for studying the occurrence, development, and treatment targets of colorectal cancer (29-31). *VIM* is present in various non-epithelial cells, particularly mesenchymal cells. Still, research shows that it is also highly expressed in colorectal cancer cells and is closely related to colorectal cancer metastasis and prognosis (32). The results showed that the *dapB* signal was almost undetectable, while the other two genes were successfully detected. Epithelial cells and mesenchymal cells were distinguished based on the specific expression of genes (Figure 2A, and the original FISH images are provided in Supplementary Fig. S2.), demonstrating that split-roll FISH can quantify gene expression levels and elucidate heterogeneous expression patterns in FFPE tumor tissue sections.

**Figure 2:**
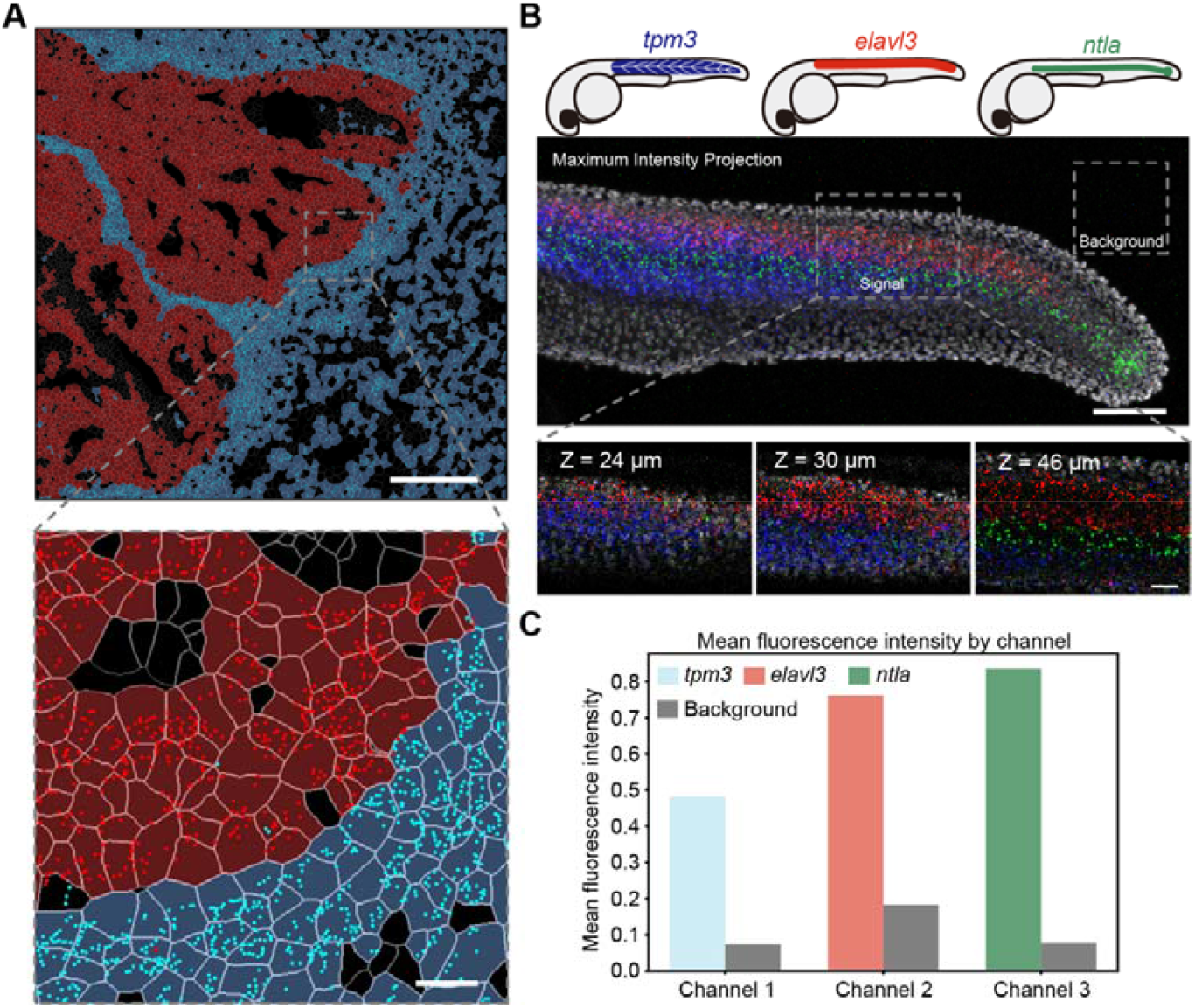
split-roll FISH detects RNAs in tissue sections and whole-mount zebrafish embryos. (**A**) Spatial visualization of epithelial cell marker *EPCAM* (red) and mesenchymal cell marker *VIM* (cyan) in colorectal cancer tissue. The colored mask shows the distribution of cells (red for epithelial cells and cyan for mesenchymal cells). The scale bar represents 200 μm. Magnified view scale bar: 20 μm. (**B**) Multiplexed mRNA expression mapping in a fixed, whole-mount zebrafish embryo. Expression profiles of three target mRNAs—*tpm3* (blue), *elavl3* (red), *ntla* (green)—are shown, with gray indicating DAPI staining. Scale bar: 100 μm. Magnified view of three planes of mRNA expression in the embryo, imaged using confocal microscopy. Scale bar: 50 μm. (**C**) Comparison of fluorescence intensity for each channel, highlighting signal intensity relative to background.

### Whole-mount zebrafish embryo *in situ* RNA detection using split-roll FISH

Achieving a high signal-to-background ratio while enabling multiplexed detection of multiple target RNAs in intact whole-mount tissues remains a significant challenge. Zebrafish serve as an outstanding model system for vertebrate developmental studies, particularly for 3D spatial gene expression analysis, enabling high-resolution visualization of gene activity patterns within intact embryos. We employed split-roll FISH to detect the expression of three genes—*tpm3* (blue), *elavl3* (red), and *ntla* (green)—in zebrafish embryos at 27 hours post-fertilization (hpf) (Figure 2B). The gene *tpm3* plays a role in skeletal myofibril assembly, *elavl3* regulates neurogenesis, and *ntla* is involved in the regulation of organ and embryonic development. All three genes have human orthologs. These results showed a high signal-to-background ratio in fixed, whole-mount zebrafish embryos (Figure 2C), demonstrating the feasibility of using split-roll FISH for whole-mount tissue gene expression analysis.

### Discrimination of SNPs by split-roll FISH

*KRAS* gene mutations are common in pancreatic ductal adenocarcinoma (PDAC) (33), non-small cell lung cancer (NSCLC) (34), and colorectal cancer (CRC) (35), and are considered to be the most common driver of human cancer (36). In cancer cells, *KRAS* gene mutations are common at codon 12, with mutations such as G12D and G12V (37). In this study, we employed split-roll FISH to identify G12V mutations in mutant (SW480) cells and wild-type (H157) cells. As shown in Figure 3A, the upstream probe of the split probe was used to locate the *KRAS* gene, while the downstream probe was used to discriminate whether a mutation occurred. Here, we termed the bases designed according to the mRNA mutation site in the probe “key bases”. We found that the second base near the 3’ end of the downstream probes of the split probe is the best “key base”, which ensures sensitivity and accuracy (Figure 3B and C, the fluorescence images for the other two base sites are shown in Supplementary Fig. S3). When the green and cyan signals appear at the same position, the signal is defined as the wild-type *KRAS* gene; and when green and red signals appear at the same position, it represents the mutant *KRAS* gene.

**Figure 3:**
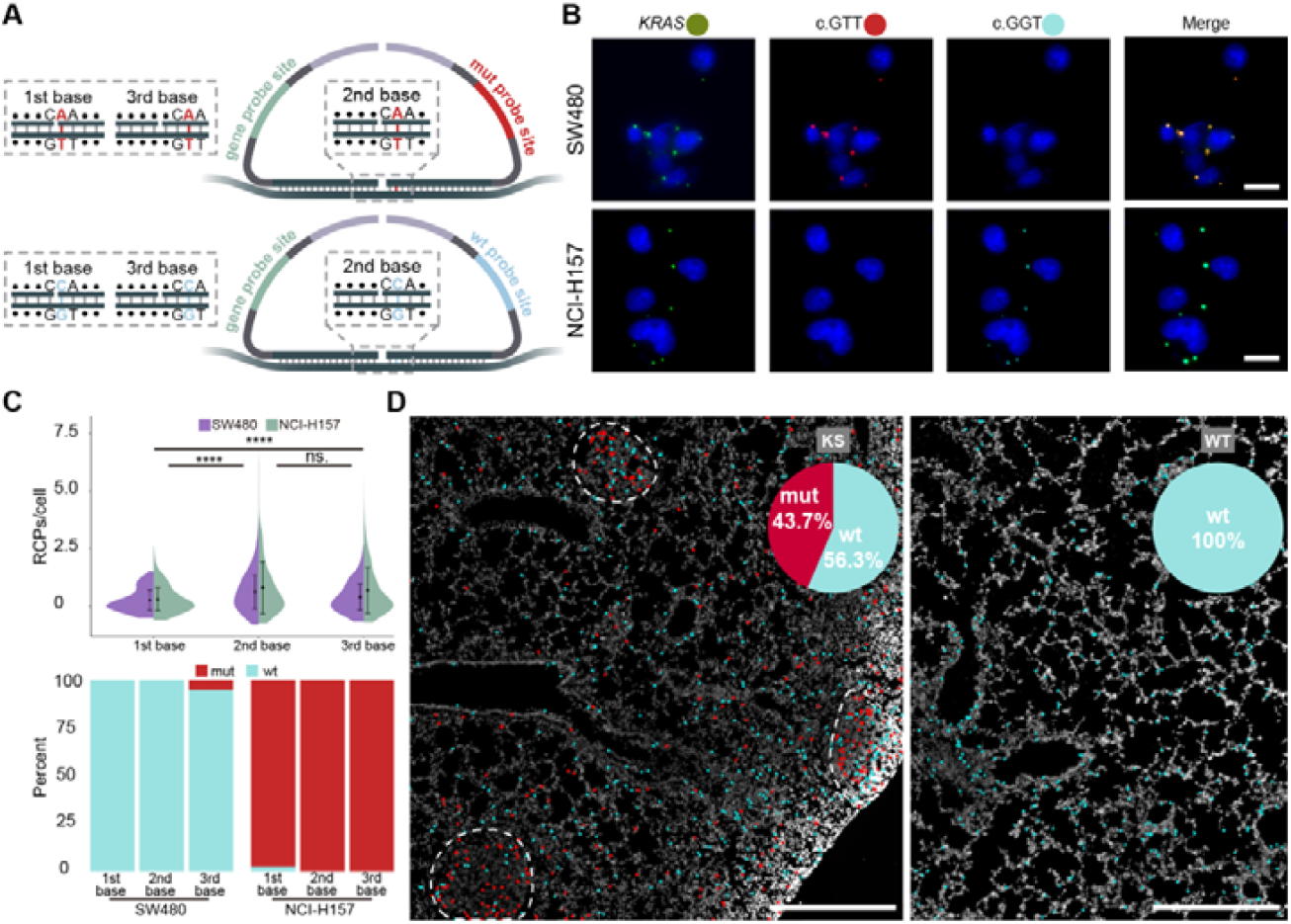
SNP detection principle, method comparison and point mutation genotype identification based on split-roll FISH. (**A**) Design principle of special split probes used in split-roll FISH for SNP detection. (**B**) Fluorescence images of identifying G12V mutations in mutant (SW480) cells and wild-type (H157) cells when the 2nd base at the 3′ end of the downstream probe is set as the key base. Scale bar: 10 μm. (**C**) Test of the position of the key base used to detect mutation sites, including comparison of the number of RCPs in single cells and the percentage of *KRAS*^G12V^ mutations in cell lines. Statistical significance between groups was assessed using one-way ANOVA. (****) indicating *P* < 0.0001. “ns” denotes no significance (*P* > 0.05). (**D**) Visualization and mutation proportion of *Kras*^G12D^ mutation in lung tissues of 9-week KS and WT mice. The white dotted circle outlines the nodules; red dots represent “mut,” cyan dots represent “wt,” and gray indicates nuclear staining with DAPI. Scale bar: 500 μm.

Next, we discriminated the *KRAS* gene for mutations in lung samples from KS and WT mice. *KRAS* with the G12D mutation was determined as mutant and designated as “mut”, while *KRAS* without the G12D mutation was determined as wild type and designated as “wt”. (Figure 3D). Currently, a variety of lung cancer models have been constructed based on *Kras*^G12D^-overexpressing mice (38). The tumors in this model are driven by *Kras*^G12D^ mutations and originate from *Sftpc*-positive type II alveolar cells. Therefore, we used the AT2 cell marker gene *Sftpc* to locate and divide the nodule area. We found that the wild-type rate of *Kras* in the lungs of WT mice was 100%, while the mutant rate of *Kras*^G12D^ in the lungs of KS mice was approximately 60%. Additionally, the mutation rate varied among different nodules. This result shows the heterogeneity of tumor development and provides a method to test the modeling effect of this model for specific tissues.

### Spatial gene expression profiling of mouse kidney by split-roll ISS

We spatially profiled renal cell types in eight mouse kidney samples across sex and disease conditions (wild-type and diabetic), revealing the complex distribution patterns of various renal cell types within the tissue (Figure 4A). Based on published single-cell transcriptome data (20), we selected marker genes for 21 mouse kidney cell types, adding differentially expressed genes identified by bulk RNA-seq between the two groups to perform ISS (Figure 4B and C). To ensure data quality, we examined the number of transcripts detected in each sample and presented the top 10 most abundant genes per sample (Supplementary Fig. S4A), which reflected consistent expression of renal marker genes (such as *Aqp1, Lrp2*) across different samples. In addition, we assessed the correlation between the number of unique genes per cell (nFeature_RNA) and the total transcript counts (nCount_RNA), revealing a strong positive correlation in all samples (Supplementary Fig. S4B), indicative of overall high-quality and reliable transcript detection.

**Figure 4:**
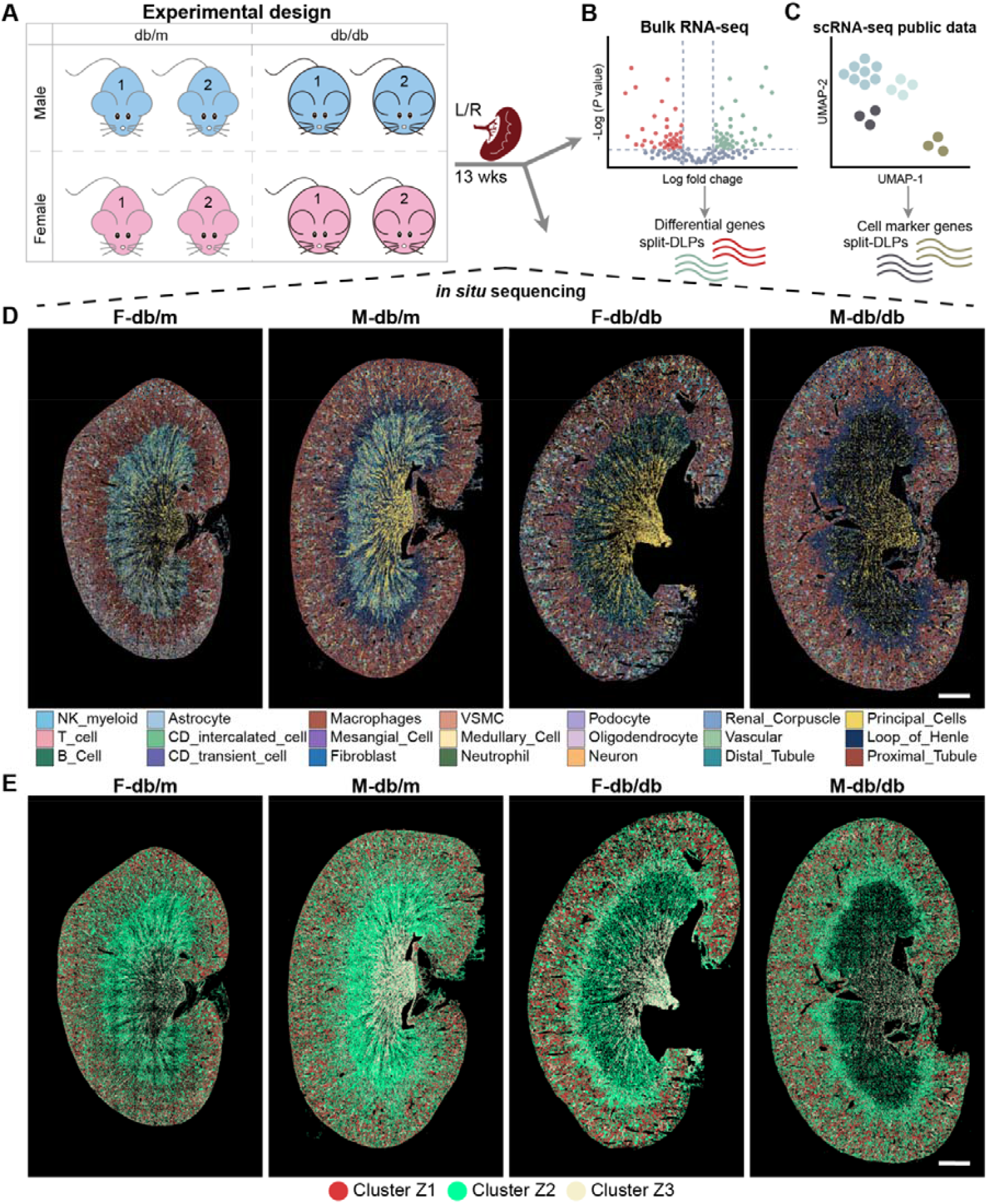
Overview of the application of split-roll ISS in analyzing cell types and gene expression spatial features in the mouse kidney of normal and diabetic mouse models. (**A**) Kidneys were harvested from 2 male and 2 female mice in each of the healthy and diabetic groups. One kidney was used for bulk RNA sequencing, while the other kidney was used for in situ sequencing. (**B**) Identifying differential genes with bulk RNA sequencing. (**C**) Identifying cell-type-specific marker genes with scRNA-seq public data. (**D**) Spatial map illustrating the distribution of cell types within kidney tissues. Scale bar: 1 mm. (**E**) Mapping kidney zonation structural labels from single-cell RNA sequencing data to spatial data revealed the spatial distribution of kidney regions. Scale bar: 1 mm.

We successfully constructed a spatial gene expression atlas of the kidneys in both healthy and diabetic mice, encompassing both male and female subjects. Specifically, we annotated and quantified the distribution of various renal cell types in different groups using nephron segment-specific markers (Supplementary Fig. S4C). Subsequently, we visualized the spatial distribution and relative abundance of all cell types (Figure 4D and Supplementary Fig. S4D). Most importantly, we used Tangram to project the single-cell RNA sequencing data of annotated kidney subregions onto the spatial transcriptomics data (Figure 4E and Supplementary Figure S5). This approach enabled us to define subregion-related spatial clusters based on the projection patterns, which we referred to as Cluster Z1, Cluster Z2, and Cluster Z3. These clusters captured the expression trends associated with subregions, such as Cluster Z1 corresponding to the cortex (Zone 1, Z1), Cluster Z2 corresponding to the outer medulla (Zone 2, Z2), and Cluster Z3 corresponding to the inner medulla (Zone 3, Z3).

In the spatial atlas of the kidneys of male and female mice, we clearly distinguished the spatial distribution of key cell types, which was consistent with the results described in the single-cell database created by Ransick et al. (20). Cell type neighborhood enrichment analysis further confirmed the rationality of the spatial distribution (Figure 5A). Notably, our atlas provides spatial validation for the existence of collecting duct transitional cells—an intermediate population between principal and intercalated cells—across the full coronal plane of the kidney at single-cell resolution (19,39). The spatial expression patterns of transitional cell markers *Insrr* and *Rhbg* closely matched their distribution in the Visium spatial transcriptomics dataset reported by Ferreira et al (40). (Supplementary Fig. S6A and B), supporting the accurate identification of this cell type. Moreover, all three collecting duct cell types are spatially enriched in their neighborhood, especially in the cluster Z1 (cortex) (Figure 5A and B, and Supplementary Fig. S7A and B). In addition, the reduction of intercalated cells and the increase of principal cells in the kidney atlas in disease states provide spatial confirmation of the view in the paper by Park et al. (19) (Supplementary Fig. S4D). More importantly, this study is the first to confirm the spatial distribution of *Aqp4* in intact mouse kidney tissue at the spatial transcriptomic level. As a key water reabsorption protein, Aqp4 is predominantly localized in the principal cells of the collecting ducts within the Z3 cluster (inner medulla), as well as in the proximal tubules at the junction of the Z1 cluster (cortex) and Z2 cluster (outer medulla)(39), highlighting the structural and functional adaptability of this protein (Figure 5C and Supplementary Fig. S7C). The tissue-context spatial localization suggests that *Aqp4*, a highly selective water channel, may fulfill distinct physiological roles in different anatomical compartments (41). Specifically, in the principal cells of the renal papillary region (Figure 5D and Supplementary Fig. S7D), *Aqp4* is highly expressed and primarily involved in mediating water transport (40,⍰141). In contrast, within the proximal tubules located in the outer stripe of the outer medulla, *Aqp4* expression is also elevated, where it likely contributes to efficient water reabsorption (40,⍰41) (Figure 5E and Supplementary Fig. S7E).

**Figure 5:**
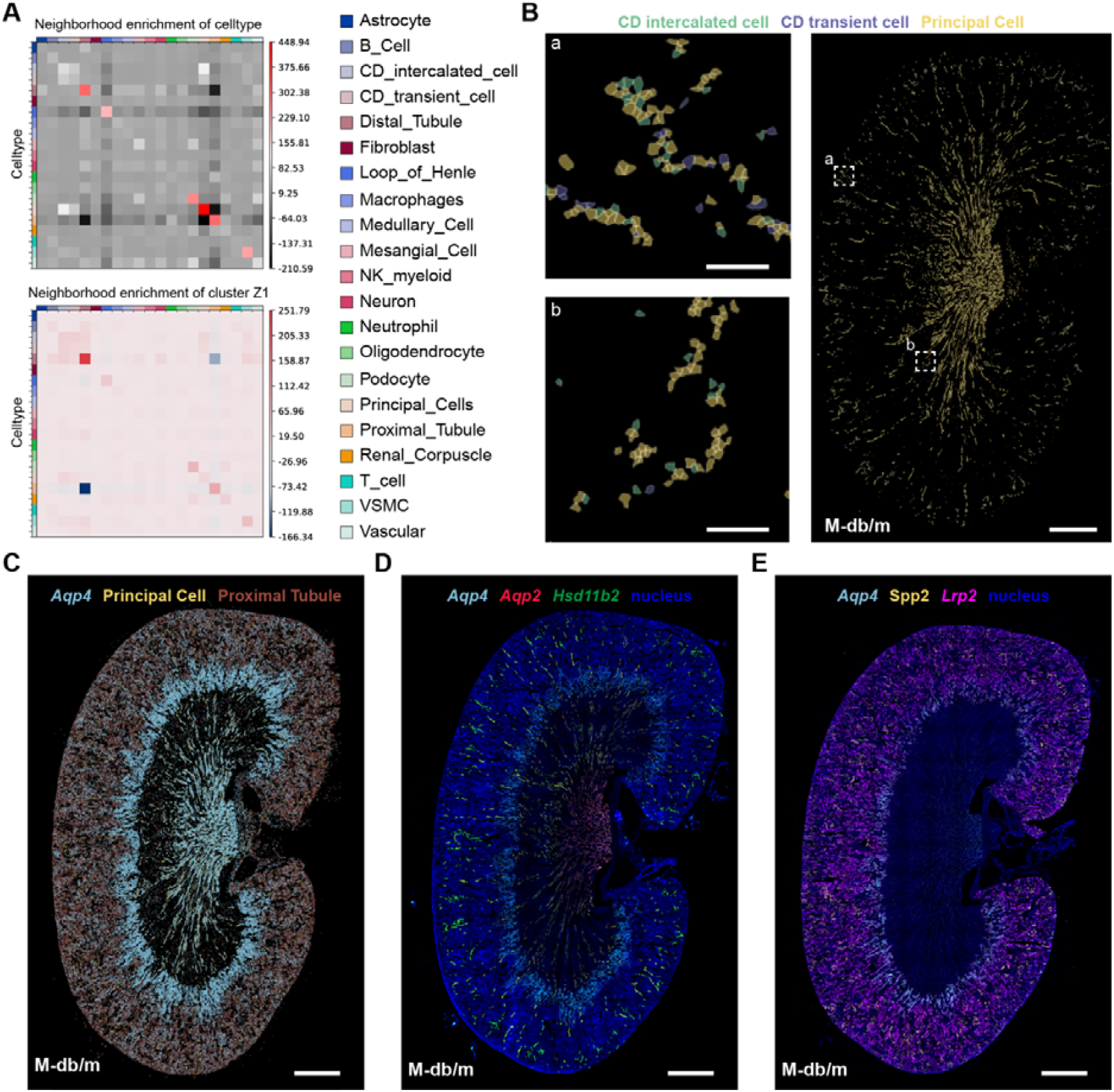
Spatial atlas of key renal cell types and functional marker expression in normal male mice. (**A**) Heatmaps of spatial neighborhood enrichment analysis showing the spatial proximity patterns among major renal cell types in the entire kidney and the cluster Z1 of normal male mice. (**B**) Spatial maps show the distribution of principal cells, intercalated cells, and transitional cells within the collecting duct. Scale bar: 1 mm. Magnified view scale bar: 100 μm. (**C**) Spatial distribution of *Aqp4* in principal cells and proximal tubules. Scale bar: 1 mm. (**D**) Spatial distribution of *Aqp4* and principal cells markers. Scale bar: 1 mm. (**E**) Spatial distribution of *Aqp4* and proximal cell markers. Scale bar: 1 mm.

### Sex-related gene differences in the kidney of diabetic mouse models

Diabetes is a complex metabolic disease. Its pathological process not only involves abnormal blood sugar regulation, but is also accompanied by physiological changes such as metabolic disorders, immune system activation, and organ damage (42,43). We performed DEG analysis on the kidneys of mice in different groups. The results showed that multiple genes involved in fatty acid metabolism (e.g., *Akrtc18*), glycogen metabolism (*Gys1*), insulin signaling pathway (*Pik3cb, Pik3ca, Map2k7* and *Cd36*), and immune response pathways (*Cd2, Cd4* and *Cd3e*) were downregulated in the diabetic group (Figure 6A). The changes in these genes may reflect the impaired metabolic function and abnormal immune function in the diabetic state, thereby aggravating the process of kidney damage (44). The results of GSVA enrichment analysis also confirmed this view (Supplementary Fig. S8A). It is worth noting that the specific expression of *Cd36* and *Ldhd* in the cortex region of the kidney supports that this region performs more metabolism-related functions, especially in maintaining water and salt balance, regulating blood pressure, acid-base balance, and other critical physiological processes (45,46). However, in the diabetic state, the kidney often undergoes metabolic reprogramming, leading to changes in fatty acid and lactate metabolic pathways (47,48), which is also suggested by the downregulation of *Cd36* and *Ldhd* expression in the cortical region of diabetic mice (Figure 6B and C, and Supplementary Fig. S8B and C).

**Figure 6:**
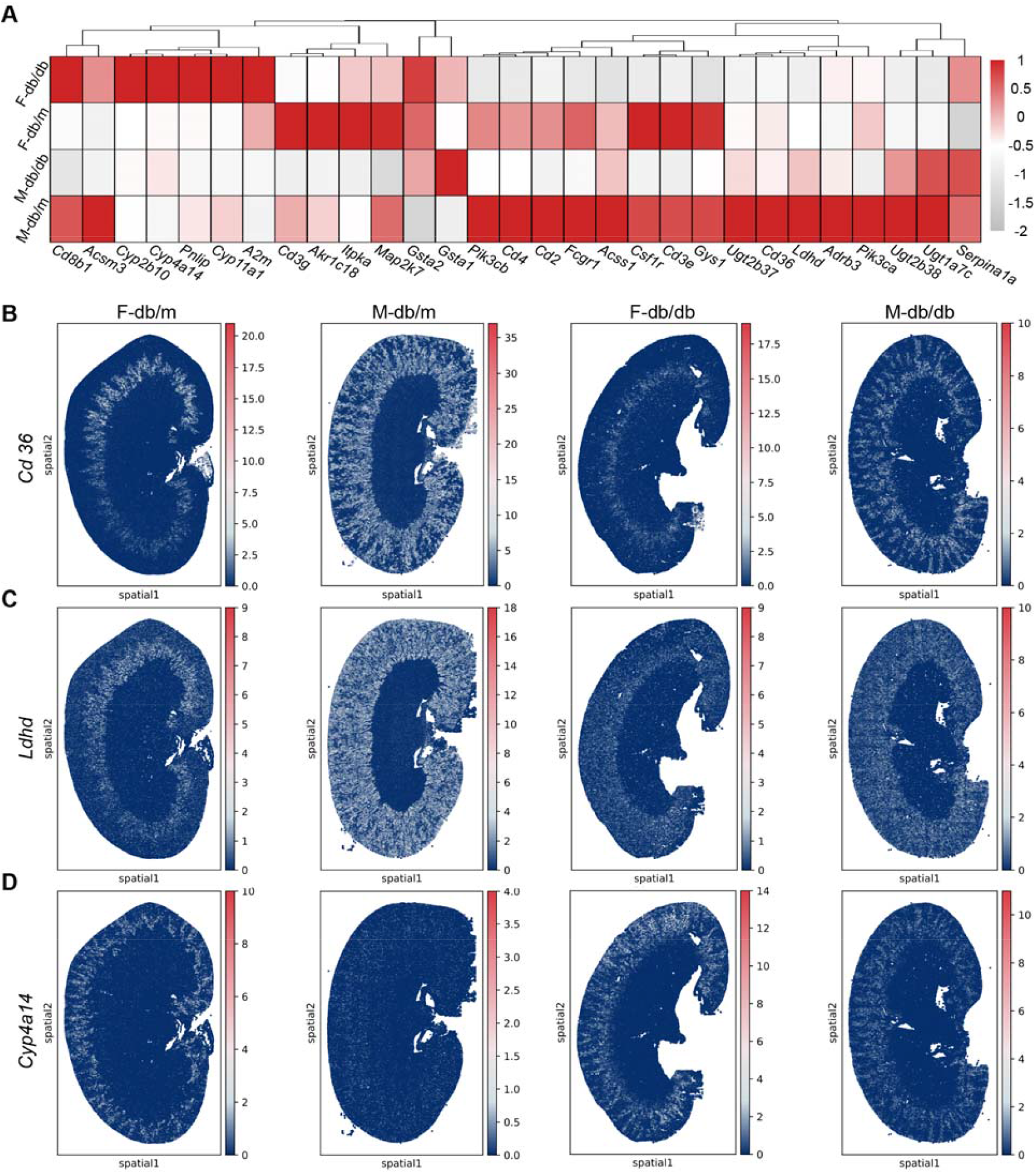
Sex-specific spatial expression changes and downregulation of key metabolic and immune-related genes in diabetic mouse kidneys. (**A**) Heat map of differentially expressed genes between the kidneys of male and female mice under normal and diabetic conditions. (**B**) Spatial distribution and expression of *Cd36* in kidney tissues under different groups. (**C**) Spatial distribution and expression of *Ldhd* in kidney tissues under different groups. (**D**) Spatial distribution and expression of *Cyp4a14* in kidney tissues under different groups.

Additionally, the transcriptome changes caused by diabetes exhibited significant gender differences. Recently, numerous studies have demonstrated significant sexual dimorphic gene expression in the proximal tubules, which is also reflected in our spatial data. However, the changes in the *Cyp4a14* gene in the diabetic group are particularly noteworthy (20,49)(Figure 6D, and Supplementary Fig. S8D). *Cyp4a14* is a gene enriched in the proximal tubules at the junction of the cortex and outer medulla of female mice, and its expression is low in normal male mice, with no obvious spatial distribution characteristics. However, the gene is highly expressed in male mice in a diseased state and exhibits a consistent spatial distribution trend similar to that of female mice. There is evidence that sex hormones may regulate changes in *Cyp4a14* expression, and that estrogen and androgen may play different roles in regulating diabetes metabolism, which may be the reason why the expression pattern of *Cyp4a14* in male diabetic mice changes to that in female mice (50-52). At the same time, the upregulated expression of this gene in diseased mice also triggered an increase in the GSVA scores of the steroid hormone biosynthesis pathway, cytochrome P450 enzyme drug metabolism pathway, and glutathione metabolism pathway in the disease group (Supplementary Fig. S8A). These results revealed changes in metabolism, inflammatory response, and oxidative stress associated with the diabetic state, providing essential clues for further research on the molecular mechanisms and structural and functional changes of diabetes.

## DISCUSSION

In this study, we successfully developed and applied a new generation of in situ RNA profiling technique, namely split-roll FISH. Our new method has significantly improved performance in many aspects compared to asmFISH, especially in detection efficiency. These are reflected not only in the optimized reaction conditions and probe design of this method, but also in its broad applicability to a wide range of sample types. First, split-roll FISH can visualize and detect single RNA molecules in human colorectal cancer tissue sections, which provides more detailed molecular information for cancer research. Secondly, split-roll FISH also performs well in whole-tissue imaging. The technology has been successfully applied to in situ RNA detection in zebrafish embryos, providing a powerful tool for developmental biology research. In addition, the technology can also distinguish SNPs and identify the genotype of point mutation mice, which shows its important application value in genetic research. By combining split-roll FISH with IISS technology, we successfully decoded the spatial distribution of 82 marker genes and created a spatial map of the kidneys in both sexes of healthy and diabetic mice. This result not only verified the distribution of *Aqp4* in the proximal tubules and collecting duct principal cells but also explored the transcriptome differences between sexes under the influence of diabetes from a spatial perspective, further helping to understand the molecular mechanism of diabetic nephropathy from the perspective of structural and functional adaptation.

In conclusion, the high sensitivity and high accuracy of split-roll FISH technology enable its broad applicability in clinical diagnostics. Notably, its effectiveness with FFPE tissue sections highlights its potential to support pathological diagnoses. Additionally, the ability of our method to identify SNPs provides a valuable tool for visualizing the spatial distribution of mutant transcripts. While split-roll FISH has demonstrated notable advantages in this study, there is still room for further optimization. Future research should focus on enhancing detection throughput and controlling costs to facilitate the broader use of this method in large-scale clinical sample analysis. Furthermore, utilizing this technology to construct high-throughput, high-resolution tissue spatial maps represents an important area for future development. Overall, split-roll FISH is a robust and versatile multiplex detection technology that offers an alternative spatial gene profiling approach for researchers.

## ACKNOWLEDGEMENTS

We would like to thank the Instrumental Analysis Center of Huaqiao University for their assistance with image acquisition and data analysis. We also want to express our sincere appreciation to Prof. Guang-Hui Jin’s group from Xiamen University for generously providing the 9-week-old LSL-*Kras*^G12D/+^; *Sftpc*-Cre mice and LSL-*Kras*^G12D/+^ mice lung FFPE tissue sections.

## AUTHOR CONTRIBUTIONS

Xueqian Xia: Data curation, Investigation, Methodology, Writing-original draft. Zhaoxiang Xie: Data curation, Investigation, Formal analysis, Visualization, Writing-original draft. Yu Yang: Data curation, Investigation, Methodology. Yanxiu Liu: Data curation, Investigation, Methodology, Writing-review & editing. Weiyan Ma: Methodology. Bixuan Zhang: Resources. Yueping Huang: Resources. Yafang Shi: Resources. Hui Lin: Resources. Lingyu Zhu: Resources. Wenhua Li: Resources. Chen Lin: Conceptualization, Funding acquisition, Supervision, Writing-review & editing. Rongqin Ke: Conceptualization, Funding acquisition, Supervision, Writing-review & editing.

## SUPPLEMENTARY DATA

Supplementary data are available at NAR online.

## DATA AVAILABILITY

The scRNA-seq data used in this study are available in the GEO database under accession number GSE129798. The Visium data used in this study are publicly available in the GEO database under accession number GES171406. Raw sequencing data, imaging data, and bulk RNA-seq data supporting the findings are available from the corresponding author upon reasonable request.

## REFERENCES

1. Gall, J.G. and Pardue, M.L. (1969) Formation and detection of RNA-DNA hybrid molecules in cytological preparations. Proc Natl Acad Sci U S A, 63, 378–383.

2. Pardue, M.L. and Gall, J.G. (1969) Molecular hybridization of radioactive DNA to the DNA of cytological preparations. Proc Natl Acad Sci U S A, 64, 600–604.

3. Femino, A.M., Fay, F.S., Fogarty, K. and Singer, R.H. (1998) Visualization of single RNA transcripts in situ. Science, 280, 585–590.

4. Raj, A., van den Bogaard, P., Rifkin, S.A., van Oudenaarden, A. and Tyagi, S. (2008) Imaging individual mRNA molecules using multiple singly labeled probes. Nat Methods, 5, 877–879.

5. Moffitt, J.R. and Zhuang, X. (2016) RNA Imaging with Multiplexed Error-Robust Fluorescence In Situ Hybridization (MERFISH). Methods Enzymol, 572, 1–49.

6. Lubeck, E., Coskun, A.F., Zhiyentayev, T., Ahmad, M. and Cai, L. (2014) Single-cell in situ RNA profiling by sequential hybridization. Nat Methods, 11, 360–361.

7. Choi, H.M., Chang, J.Y., Trinh le, A., Padilla, J.E., Fraser, S.E. and Pierce, N.A. (2010) Programmable in situ amplification for multiplexed imaging of mRNA expression. Nat Biotechnol, 28, 1208–1212.

8. Wang, F., Flanagan, J., Su, N., Wang, L.C., Bui, S., Nielson, A., Wu, X., Vo, H.T., Ma, X.J. and Luo, Y. (2012) RNAscope: a novel in situ RNA analysis platform for formalin-fixed, paraffin-embedded tissues. J Mol Diagn, 14, 22–29.

9. Larsson, C., Grundberg, I., Söderberg, O. and Nilsson, M. (2010) In situ detection and genotyping of individual mRNA molecules. Nat Methods, 7, 395–397.

10. Sountoulidis, A., Liontos, A., Nguyen, H.P., Firsova, A.B., Fysikopoulos, A., Qian, X., Seeger, W., Sundström, E., Nilsson, M. and Samakovlis, C. (2020) SCRINSHOT enables spatial mapping of cell states in tissue sections with single-cell resolution. PLoS Biol, 18, e3000675.

11. Ke, R., Mignardi, M., Pacureanu, A., Svedlund, J., Botling, J., Wählby, C. and Nilsson, M. (2013) In situ sequencing for RNA analysis in preserved tissue and cells. Nat Methods, 10, 857–860.

12. Wang, X., Allen, W.E., Wright, M.A., Sylwestrak, E.L., Samusik, N., Vesuna, S., Evans, K., Liu, C., Ramakrishnan, C., Liu, J. et al. (2018) Three-dimensional intact-tissue sequencing of single-cell transcriptional states. Science, 361.

13. Soares, R.R.G., Madaboosi, N. and Nilsson, M. (2021) Rolling Circle Amplification in Integrated Microsystems: An Uncut Gem toward Massively Multiplexed Pathogen Diagnostics and Genotyping. Acc Chem Res, 54, 3979–3990.

14. Lin, C., Jiang, M., Liu, L., Chen, X., Zhao, Y., Chen, L., Hong, Y., Wang, X., Hong, C., Yao, X. et al. (2021) Imaging of individual transcripts by amplification-based single-molecule fluorescence in situ hybridization. N Biotechnol, 61, 116–123.

15. Fredriksson, S., Gullberg, M., Jarvius, J., Olsson, C., Pietras, K., Gústafsdóttir, S.M., Ostman, A. and Landegren, U. (2002) Protein detection using proximity-dependent DNA ligation assays. Nat Biotechnol, 20, 473–477.

16. Ho, C.K., Van Etten, J.L. and Shuman, S. (1997) Characterization of an ATP-dependent DNA ligase encoded by Chlorella virus PBCV-1. J Virol, 71, 1931–1937.

17. Tang, X., Chen, J., Zhang, X., Liu, X., Xie, Z., Wei, K., Qiu, J., Ma, W., Lin, C. and Ke, R. (2023) Improved in situ sequencing for high-resolution targeted spatial transcriptomic analysis in tissue sections. J Genet Genomics, 50, 652–660.

18. Choi, H.M., Beck, V.A. and Pierce, N.A. (2014) Next-generation in situ hybridization chain reaction: higher gain, lower cost, greater durability. ACS Nano, 8, 4284–4294.

19. Park, J., Shrestha, R., Qiu, C., Kondo, A., Huang, S., Werth, M., Li, M., Barasch, J. and Suszták, K. (2018) Single-cell transcriptomics of the mouse kidney reveals potential cellular targets of kidney disease. Science, 360, 758–763.

20. Ransick, A., Lindström, N.O., Liu, J., Zhu, Q., Guo, J.J., Alvarado, G.F., Kim, A.D., Black, H.G., Kim, J. and McMahon, A.P. (2019) Single-Cell Profiling Reveals Sex, Lineage, and Regional Diversity in the Mouse Kidney. Dev Cell, 51, 399-413.e397.

21. Qiu, H., Jin, B.M., Wang, Z.F., Xu, B., Zheng, Q.F., Zhang, L., Zhu, L.Y., Shi, S., Yuan, J.B., Lin, X. et al. (2020) MEN1 deficiency leads to neuroendocrine differentiation of lung cancer and disrupts the DNA damage response. Nat Commun, 11, 1009.

22. van der Walt, S., Schönberger, J.L., Nunez-Iglesias, J., Boulogne, F., Warner, J.D., Yager, N., Gouillart, E. and Yu, T. (2014) scikit-image: image processing in Python. PeerJ, 2, e453.

23. Stringer, C. and Pachitariu, M. (2025) Cellpose3: one-click image restoration for improved cellular segmentation. Nat Methods, 22, 592–599.

24. Biancalani, T., Scalia, G., Buffoni, L., Avasthi, R., Lu, Z., Sanger, A., Tokcan, N., Vanderburg, C.R., Segerstolpe, Å., Zhang, M. et al. (2021) Deep learning and alignment of spatially resolved single-cell transcriptomes with Tangram. Nat Methods, 18, 1352–1362.

25. Palla, G., Spitzer, H., Klein, M., Fischer, D., Schaar, A.C., Kuemmerle, L.B., Rybakov, S., Ibarra, I.L., Holmberg, O., Virshup, I. et al. (2022) Squidpy: a scalable framework for spatial omics analysis. Nat Methods, 19, 171–178.

26. Hao, Y., Hao, S., Andersen-Nissen, E., Mauck, W.M., 3rd, Zheng, S., Butler, A., Lee, M.J., Wilk, A.J., Darby, C., Zager, M. et al. (2021) Integrated analysis of multimodal single-cell data. Cell, 184, 3573-3587.e3529.

27. Hänzelmann, S., Castelo, R. and Guinney, J. (2013) GSVA: gene set variation analysis for microarray and RNA-seq data. BMC Bioinformatics, 14, 7.

28. Yu, G., Wang, L.G., Han, Y. and He, Q.Y. (2012) clusterProfiler: an R package for comparing biological themes among gene clusters. Omics, 16, 284–287.

29. Herlyn, M., Steplewski, Z., Herlyn, D. and Koprowski, H. (1979) Colorectal carcinoma-specific antigen: detection by means of monoclonal antibodies. Proc Natl Acad Sci U S A, 76, 1438–1442.

30. Osta, W.A., Chen, Y., Mikhitarian, K., Mitas, M., Salem, M., Hannun, Y.A., Cole, D.J. and Gillanders, W.E. (2004) EpCAM is overexpressed in breast cancer and is a potential target for breast cancer gene therapy. Cancer Res, 64, 5818–5824.

31. Keller, L., Werner, S. and Pantel, K. (2019) Biology and clinical relevance of EpCAM. Cell Stress, 3, 165–180.

32. 32. Wang, Q., Zhu, G., Lin, C., Lin, P., Chen, H., He, R., Huang, Y., Yang, S. and Ye, J. (2021) Vimentin affects colorectal cancer proliferation, invasion, and migration via regulated by activator protein 1. J Cell Physiol, 236, 7591–7604.

33. Gurreri, E., Genovese, G., Perelli, L., Agostini, A., Piro, G., Carbone, C. and Tortora, G. (2023) KRAS-Dependency in Pancreatic Ductal Adenocarcinoma: Mechanisms of Escaping in Resistance to KRAS Inhibitors and Perspectives of Therapy. Int J Mol Sci, 24.

34. Cascetta, P., Marinello, A., Lazzari, C., Gregorc, V., Planchard, D., Bianco, R., Normanno, N. and Morabito, A. (2022) KRAS in NSCLC: State of the Art and Future Perspectives. Cancers (Basel), 14.

35. Zhu, G., Pei, L., Xia, H., Tang, Q. and Bi, F. (2021) Role of oncogenic KRAS in the prognosis, diagnosis and treatment of colorectal cancer. Mol Cancer, 20, 143.

36. Prior, I.A., Lewis, P.D. and Mattos, C. (2012) A comprehensive survey of Ras mutations in cancer. Cancer Res, 72, 2457–2467.

37. Mondal, K., Posa, M.K., Shenoy, R.P. and Roychoudhury, S. (2024) KRAS Mutation Subtypes and Their Association with Other Driver Mutations in Oncogenic Pathways. Cells, 13.

38. Kwon, M.C. and Berns, A. (2013) Mouse models for lung cancer. Mol Oncol, 7, 165–177.

39. Novella-Rausell, C., Grudniewska, M., Peters, D.J.M. and Mahfouz, A. (2023) A comprehensive mouse kidney atlas enables rare cell population characterization and robust marker discovery. iScience, 26, 106877.

40. Melo Ferreira, R., Sabo, A.R., Winfree, S., Collins, K.S., Janosevic, D., Gulbronson, C.J., Cheng, Y.H., Casbon, L., Barwinska, D., Ferkowicz, M.J. et al. (2021) Integration of spatial and single-cell transcriptomics localizes epithelial cell-immune cross-talk in kidney injury. JCI Insight, 6.

41. van Hoek, A.N., Ma, T., Yang, B., Verkman, A.S. and Brown, D. (2000) Aquaporin-4 is expressed in basolateral membranes of proximal tubule S3 segments in mouse kidney. Am J Physiol Renal Physiol, 278, F310–316.

42. Dilworth, L., Facey, A. and Omoruyi, F. (2021) Diabetes Mellitus and Its Metabolic Complications: The Role of Adipose Tissues. Int J Mol Sci, 22.

43. Tsalamandris, S., Antonopoulos, A.S., Oikonomou, E., Papamikroulis, G.A., Vogiatzi, G., Papaioannou, S., Deftereos, S. and Tousoulis, D. (2019) The Role of Inflammation in Diabetes: Current Concepts and Future Perspectives. Eur Cardiol, 14, 50–59.

44. Amorim, R.G., Guedes, G.D.S., Vasconcelos, S.M.L. and Santos, J.C.F. (2019) Kidney Disease in Diabetes Mellitus: Cross-Linking between Hyperglycemia, Redox Imbalance and Inflammation. Arq Bras Cardiol, 112, 577–587.

45. Dumas, S.J., Meta, E., Borri, M., Luo, Y., Li, X., Rabelink, T.J. and Carmeliet, P. (2021) Phenotypic diversity and metabolic specialization of renal endothelial cells. Nat Rev Nephrol, 17, 441–464.

46. Gray, L.R., Tompkins, S.C. and Taylor, E.B. (2014) Regulation of pyruvate metabolism and human disease. Cell Mol Life Sci, 71, 2577–2604.

47. Wang, M., Pang, Y., Guo, Y., Tian, L., Liu, Y., Shen, C., Liu, M., Meng, Y., Cai, Z., Wang, Y. et al. (2022) Metabolic reprogramming: A novel therapeutic target in diabetic kidney disease. Front Pharmacol, 13, 970601.

48. Niu, H., Ren, X., Tan, E., Wan, X., Wang, Y., Shi, H., Hou, Y. and Wang, L. (2023) CD36 deletion ameliorates diabetic kidney disease by restoring fatty acid oxidation and improving mitochondrial function. Ren Fail, 45, 2292753.

49. Wu, H., Lai, C.F., Chang-Panesso, M. and Humphreys, B.D. (2020) Proximal Tubule Translational Profiling during Kidney Fibrosis Reveals Proinflammatory and Long Noncoding RNA Expression Patterns with Sexual Dimorphism. J Am Soc Nephrol, 31, 23–38.

50. Lei, Z., Wu, H., Yang, Y., Hu, Q., Lei, Y., Liu, W., Nie, Y., Yang, L., Zhang, X., Yang, C. et al. (2021) Ovariectomy Impaired Hepatic Glucose and Lipid Homeostasis and Altered the Gut Microbiota in Mice With Different Diets. Front Endocrinol (Lausanne), 12, 708838.

51. Zhang, Y. and Klaassen, C.D. (2013) Hormonal regulation of Cyp4a isoforms in mouse liver and kidney. Xenobiotica, 43, 1055–1063.

52. Mauvais-Jarvis, F., Clegg, D.J. and Hevener, A.L. (2013) The role of estrogens in control of energy balance and glucose homeostasis. Endocr Rev, 34, 309–338.

