## Supplementary Data for "Efficient spatial gene expression profiling using split-probe ligation and rolling circle amplification"

Table of contents

Fig. S1. Optimization of conditions for split-probe hybridization and circularization.

Fig. S2. *VIM* and *EPCAM* genes were detected in human colorectal cancer samples using split-roll FISH.

Fig. S3. The detection results of the bases at the first and third positions near the 3′ end of the downstream probes of the split probe are used as the bases for identifying SNPs.

Fig. S4. Quality control and cell type characterization in the spatial transcriptomic atlas of mouse kidneys.

Fig. S5. Spatial mapping of cell types and zonation in additional kidney samples.

Fig. S6. Spatial expression validation of transitional cell markers *Insrr* and *Rhbg* in the collecting duct system of the mouse kidney.

Fig. S7. Spatial atlas of key renal cell types and functional marker expression in normal female mice.

Fig. S8. Sex-specific spatial transcriptomic signatures and validation in additional kidney samples.

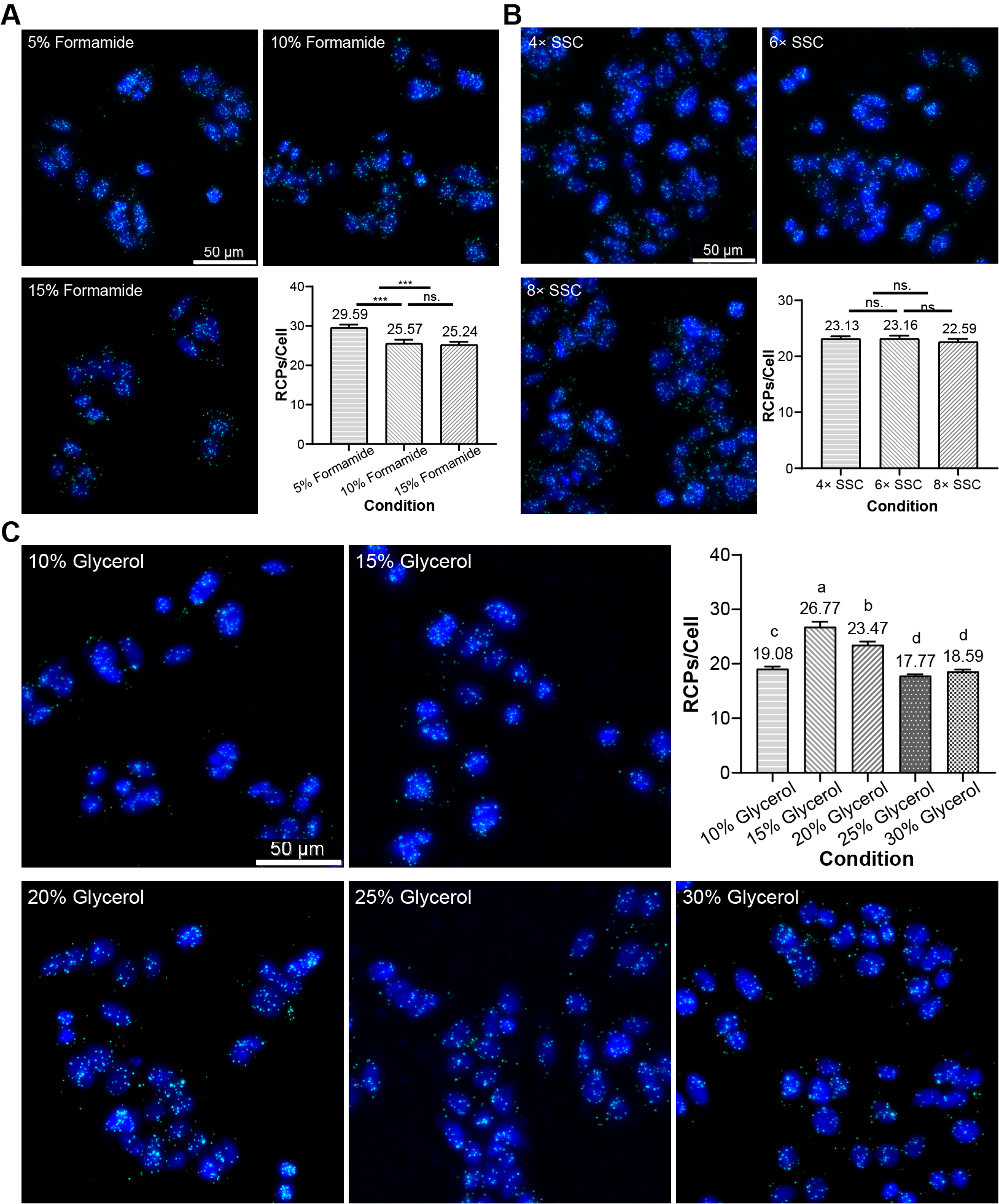

**Fig. S1.** Optimization of split-probe hybridization and circularization conditions by comparing *KRAS* (cyan) detection rates in single SW480 cells. (**A**) To determine the effect of formamide concentration on split-probe hybridization efficiency, gradient formamide concentrations were tested, and 5% formamide was selected as the optimal concentration in the split-probe hybridization buffer, as it yielded the highest RCPs per cell. (**B**) To evaluate the effect of gradient SSC concentrations on split-probe hybridization efficiency, no statistically significant differences were observed in RCPs per cell among 4×, 5×, and 6× SSC. Consequently, the original 6× SSC was retained as the concentration in the split−probe hybridization buffer. Data are presented as mean ± SEM. One-way ANOVA was performed, with (***) indicating *P* < 0.001. "ns." denotes no significance (*P* > 0.05). (**C**) The effect of glycerol concentration on split-probe circularization efficiency was assessed using gradient glycerol concentrations. 15% glycerol was selected as the optimal concentration in the circularization reaction buffer, as it provided the highest RCPs per cell. One-way ANOVA, groups labeled with the same letter did not differ significantly (*P* > 0.05), whereas groups with different letters showed significant differences (Duncan′s test).

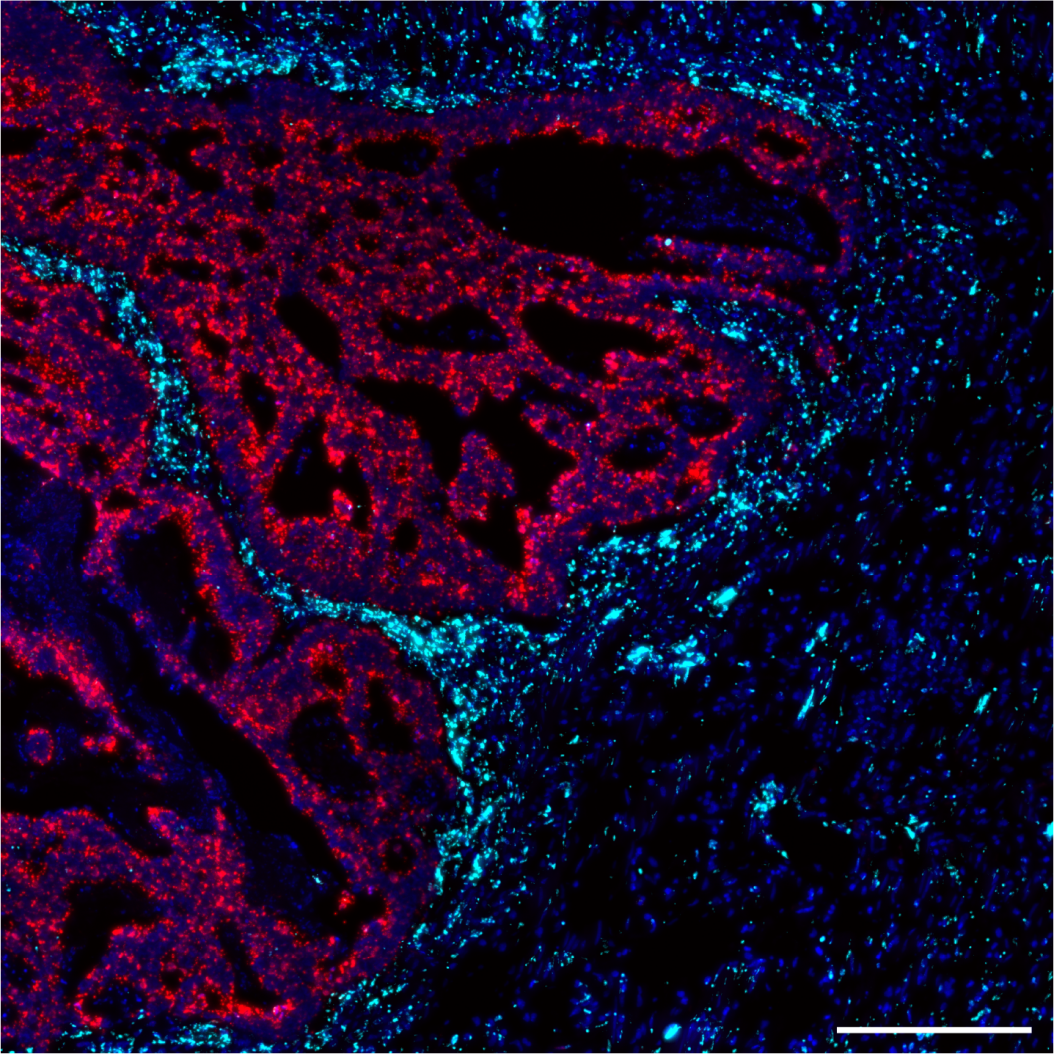

**Fig. S2.** *VIM* and *EPCAM* genes were detected in human colorectal cancer samples using split-roll FISH. Cyan signals correspond to *VIM*, red signals correspond to *EPCAM*, and the scale bar is 200 μm.

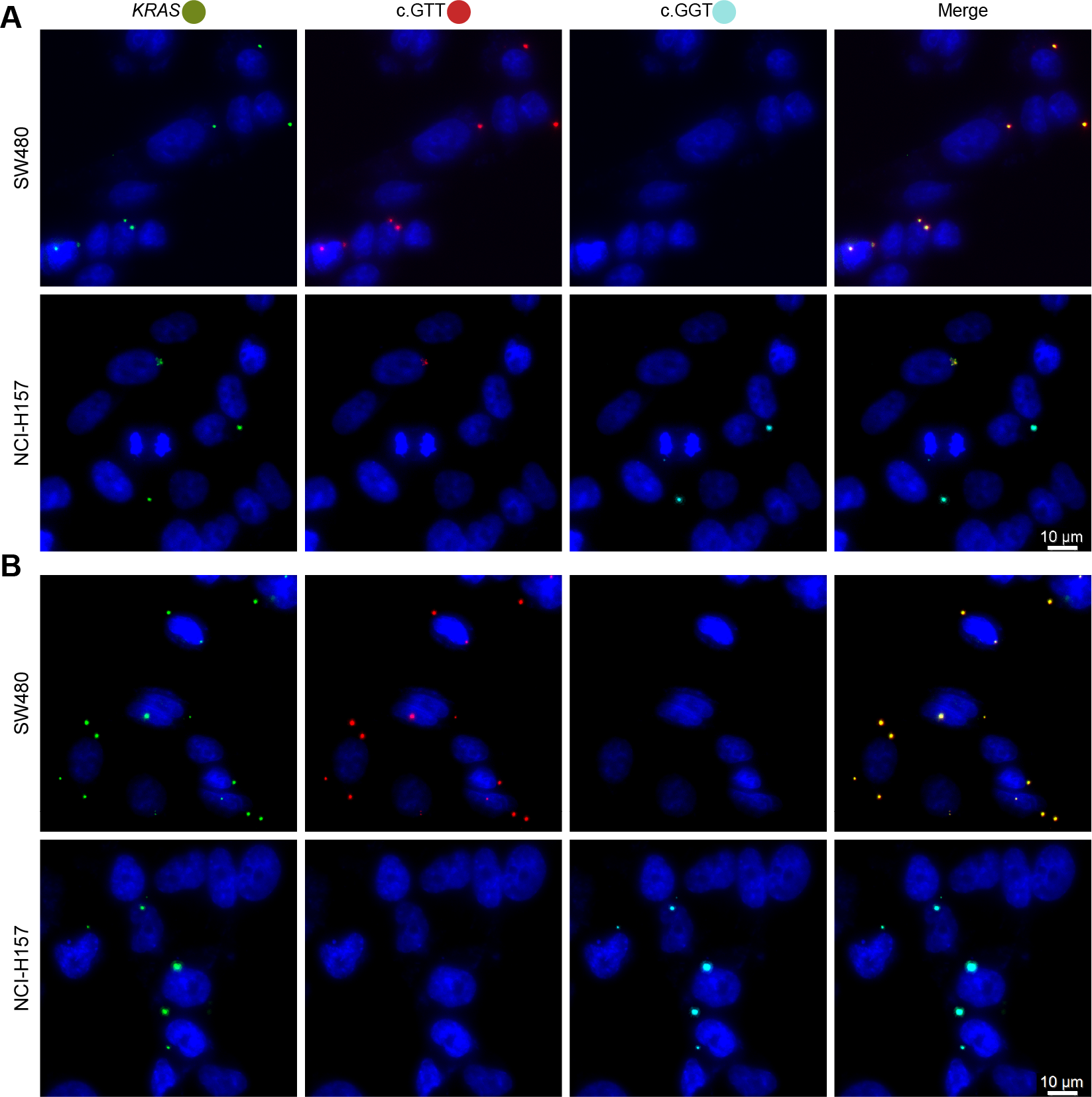

**Fig. S3.** The detection results of the discrimination bases at the first and third positions near the 3′ end of the downstream probes of the split probe are used as the bases for identifying SNPs. (**A**) The results of discrimination bases set at the first position near the 3′ end of the downstream probes of the split probe. (**B**) The results of the discrimination bases set at the third position near the 3′ end of the downstream probes of the split probe. Scale bars, 10 μm.

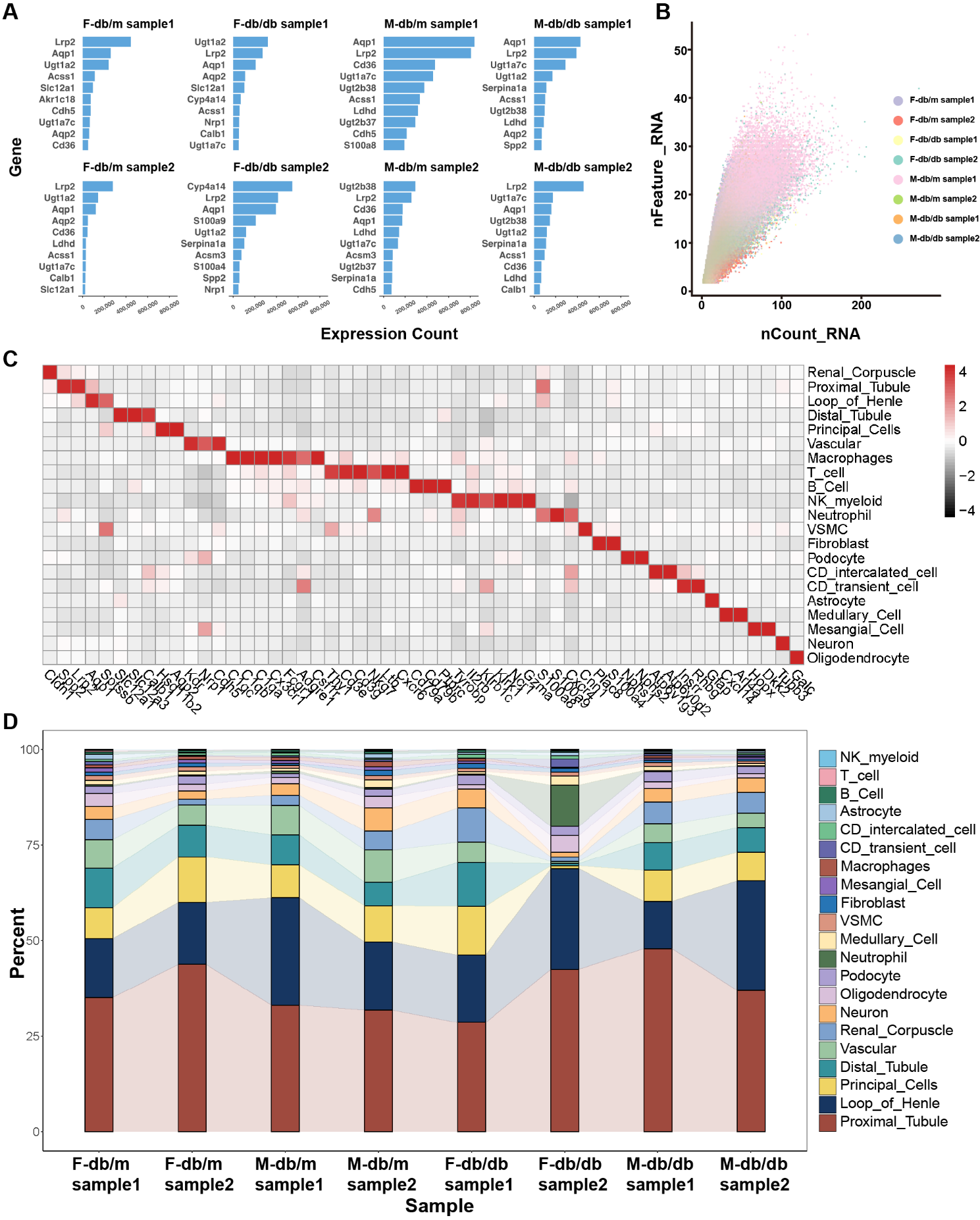

**Fig. S4.** Quality control and cell type characterization in the spatial transcriptomic atlas of mouse kidneys. (**A**) Bar plots showing the top 10 most abundantly expressed genes detected in each sample across different conditions (F-db/m, F-db/db, M-db/m, M-db/db). Gene expression levels are represented by total expression counts. (**B**) Scatter plot of the number of unique genes detected per spot (nFeature_RNA) versus total UMI counts (nCount_RNA) for each sample, illustrating quality and consistency across all eight datasets. Each dot represents a cell, and samples are color-coded by group. (**C**) The heatmap of cell type marker expression in kidney tissues, showing the expression patterns of different cell type markers. (**D**) Stacked bar chart representing the relative proportions of each cell type in different sample.

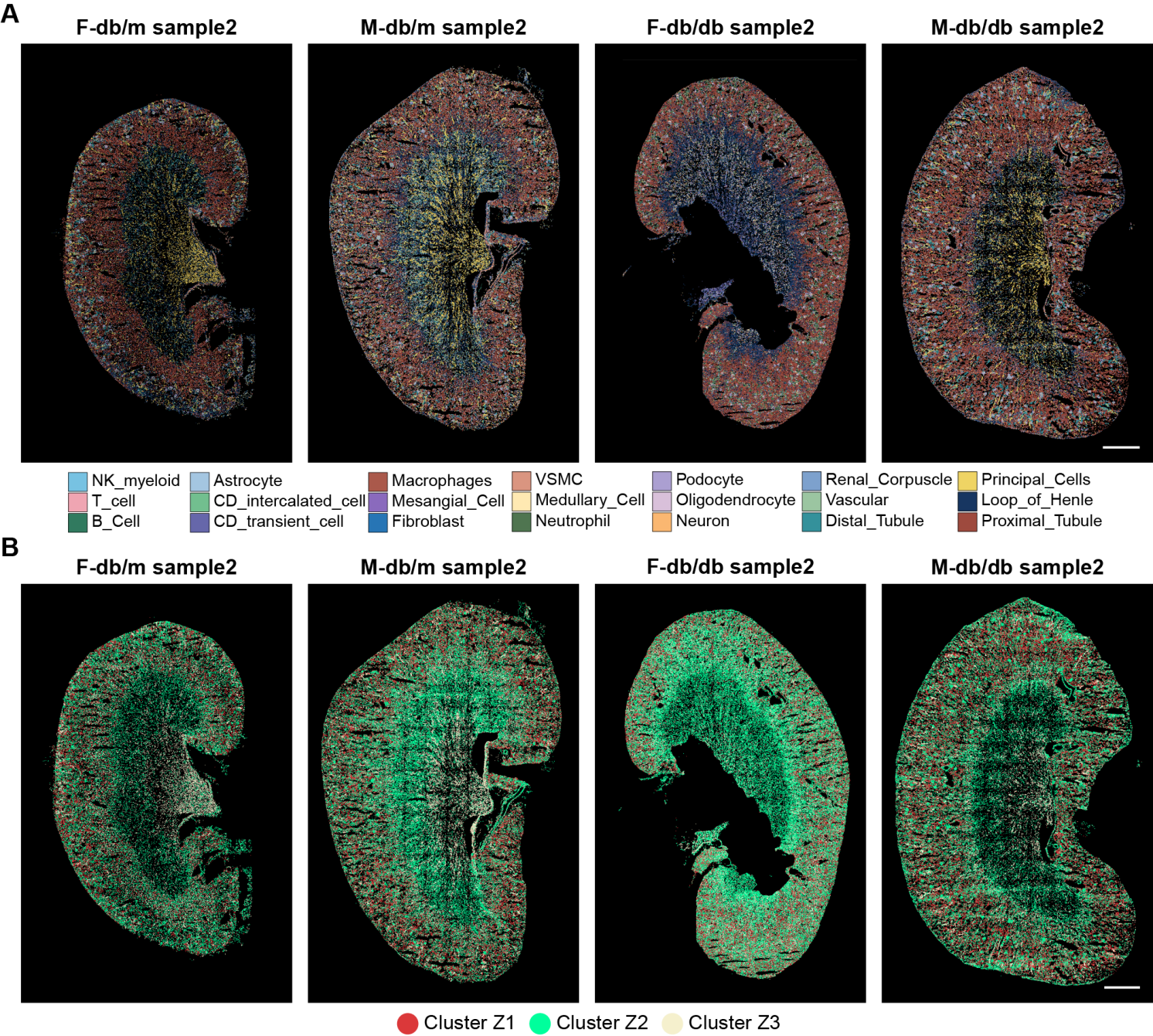

**Fig. S5.** Spatial mapping of cell types and zonation in additional kidney samples. (**A**) Spatial map illustrating the distribution of cell types within kidney tissues from additional mouse samples. The annotation was performed using cell type-specific marker genes. Scale bar: 1 mm. (**B**) Integration of kidney zonation structural labels from single-cell RNA sequencing data with spatial transcriptomics of additional samples revealed the spatial distribution of kidney regions. Scale bar: 1 mm.

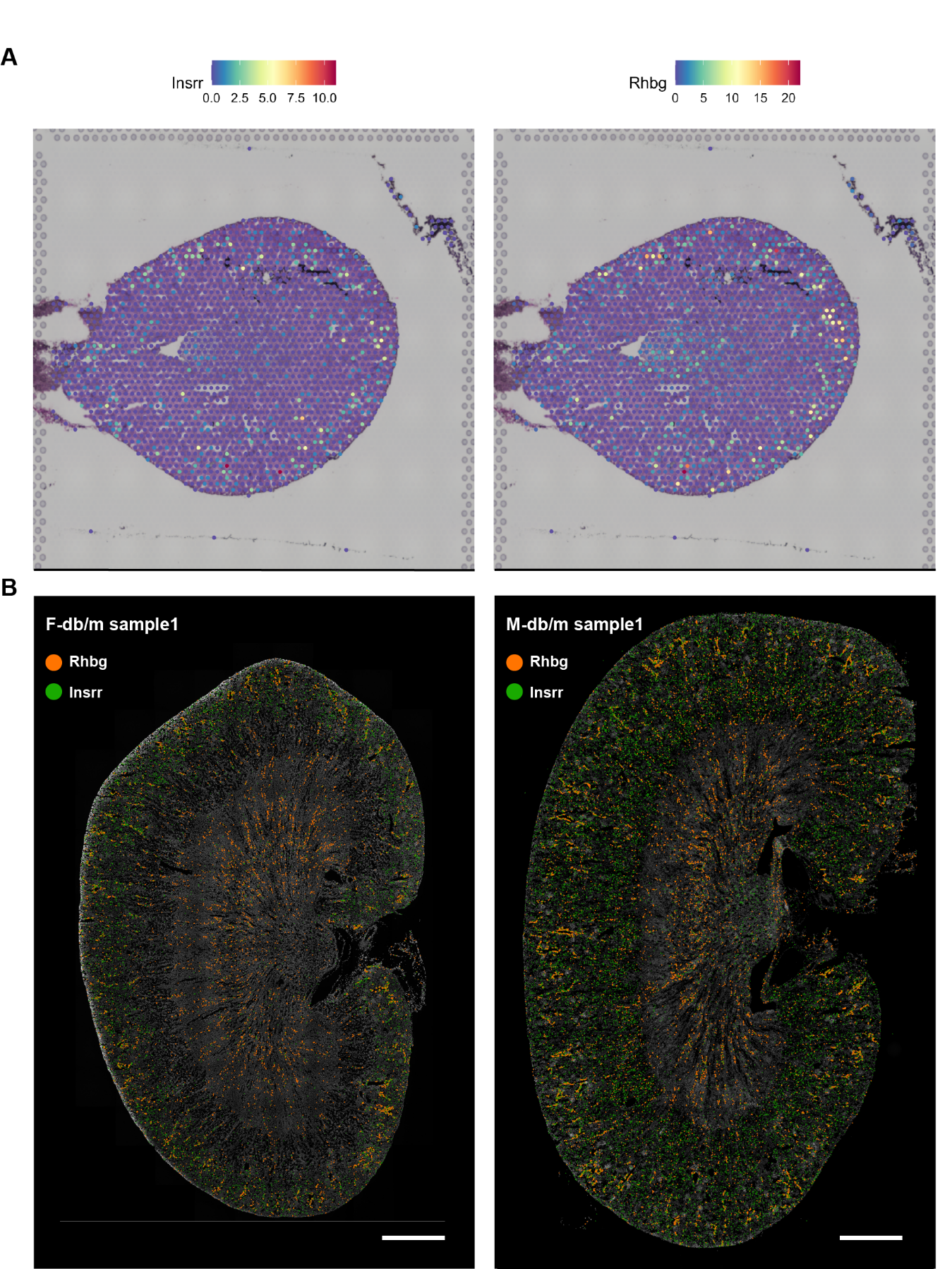

**Fig. S6.** Spatial expression validation of transitional cell markers *Insrr* and *Rhbg* in the collecting duct system of the mouse kidney. (**A**) Spatial feature plots from the Visium dataset show the expression of Insrr (left) and Rhbg (right) in mouse kidney sections. Cooler colors indicate lower expression levels, while warmer colors represent higher expression. (**B**) Spatial transcriptomic maps of *Insrr* (green) and *Rhbg* (orange) expression in representative female (left) and male (right) mouse kidney sections. Scale bar: 1 mm.

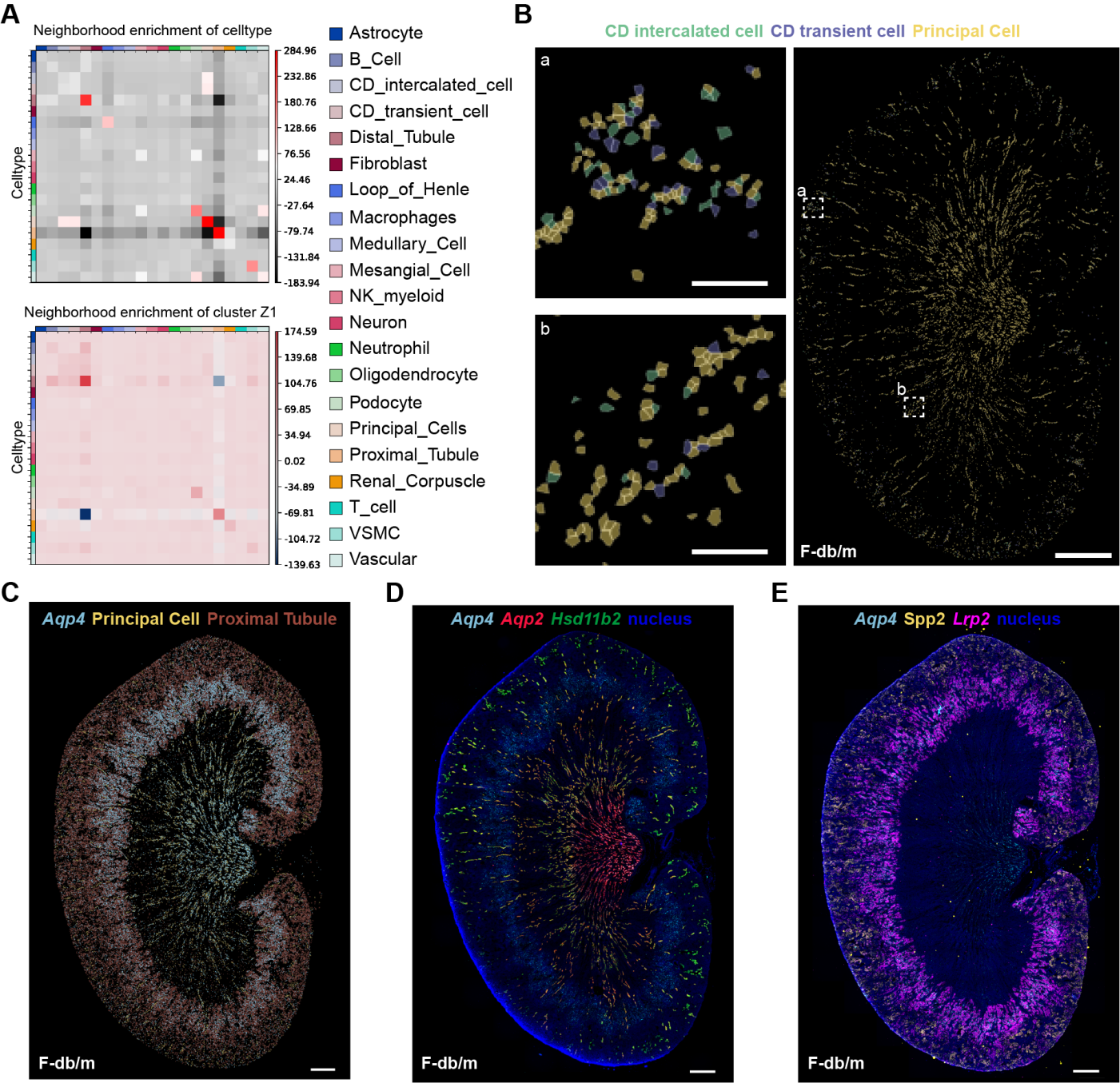

**Fig. S7.** Spatial atlas of key renal cell types and functional marker expression in normal female mice. (**A**) Heatmaps of spatial neighborhood enrichment analysis showing the spatial proximity patterns among major renal cell types in the entire kidney and the cluster Z1 of normal female mice. (**B**) Spatial maps show the distribution of principal cells, intercalated cells, and transitional cells within the collecting duct. Scale bar: 1 mm. Magnified view scale bar: 100 μm. (**C**) Spatial distribution of *Aqp4* in principal cells and proximal tubules. Scale bar: 1 mm. (**D**) Spatial distribution of *Aqp4* and principal cell markers. Scale bar: 1 mm. (**E**) Spatial distribution of *Aqp4* and proximal tubule cell markers. Scale bar: 1 mm.

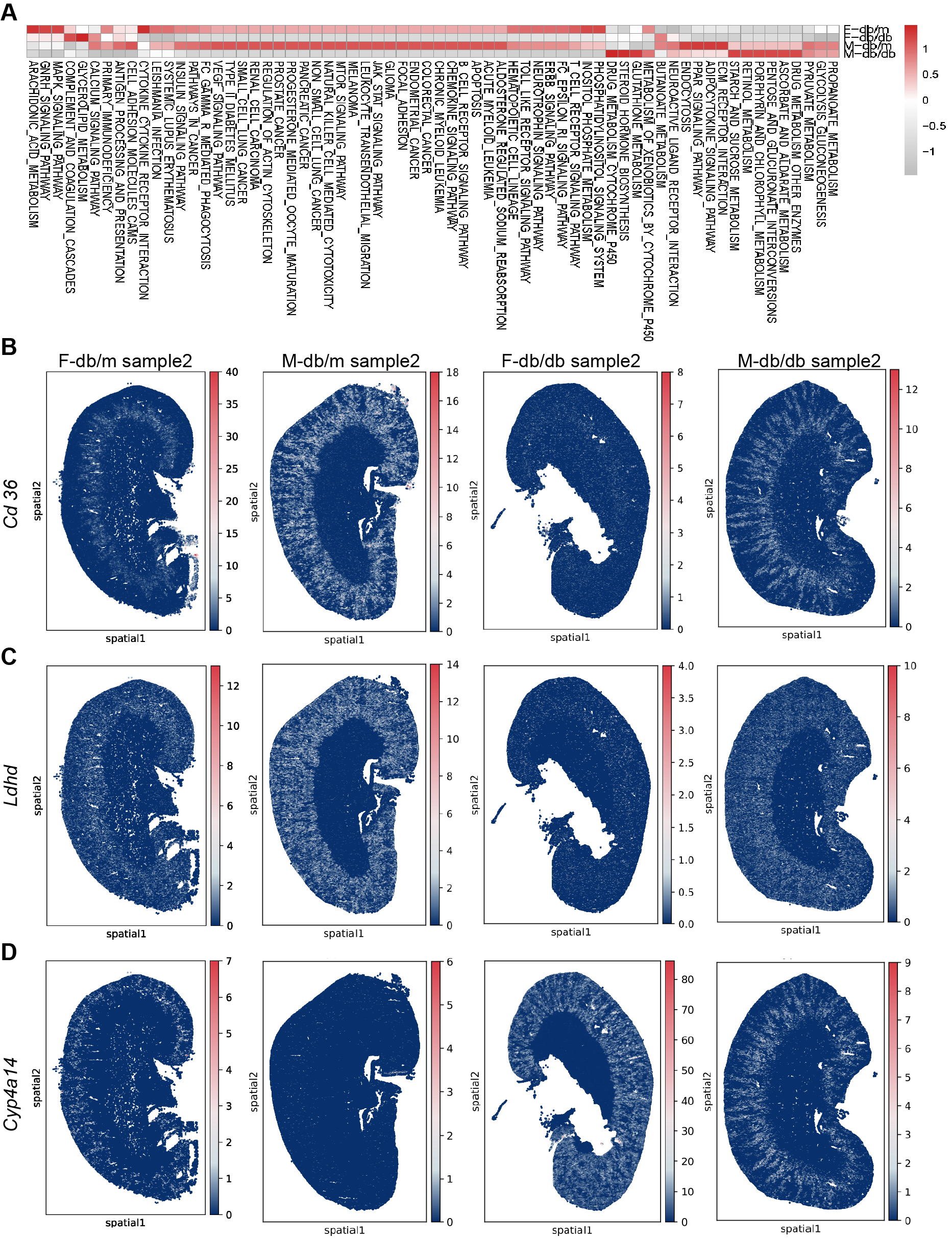

**Fig. S8.** Sex-specific spatial transcriptomic signatures and validation in additional kidney samples. (**A**) Heatmap showing the GSVA enrichment scores of selected KEGG pathways across normal and diabetic groups in male and female mice. (**B**) Spatial distribution and expression levels of *Cd36* in kidney tissues from additional normal and diabetic samples. (**C**) Spatial distribution and expression levels of *Ldhd* in kidney tissues from additional normal and diabetic samples. (**D**) Spatial distribution and expression levels of *Cyp4a14* in kidney tissues from additional normal and diabetic samples.
